# A membrane-associated condensate drives paternal epigenetic inheritance in *C. elegans*

**DOI:** 10.1101/2020.12.10.417311

**Authors:** Jan Schreier, Sabrina Dietz, Antonio M. de Jesus Domingues, Ann-Sophie Seistrup, Dieu An H. Nguyen, Elizabeth J. Gleason, Huiping Ling, Steven W. L’Hernault, Carolyn M. Phillips, Falk Butter, René F. Ketting

## Abstract

Transgenerational epigenetic inheritance (TEI) describes the transmission of gene-regulatory information across generations without altering DNA sequences, and allows priming of offspring towards transposable elements (TEs) and changing environmental conditions. One important mechanism that acts in TEI is based on small non-coding RNAs. Whereas factors for maternal inheritance of small RNAs have been identified, paternal inheritance is poorly understood, as much of the cellular content is extruded during spermatogenesis. We identify a phase separation-based mechanism, driven by the protein PEI-1, which is characterized by a BTB-BACK domain and an intrinsically disordered region (IDR). PEI-1 specifically secures the Argonaute protein WAGO-3 within maturing sperm in *C. elegans*. Localization of PEI granules in mature sperm is coupled, via S-palmitoylation, to myosin-driven transport of membranous organelles. *pei-1*-like genes are also found in human and often expressed in testis, suggesting that the here identified mechanism may be broadly conserved.

## INTRODUCTION

Transgenerational epigenetic inheritance (TEI) is a non-genetic way of inheriting gene regulatory information across generations, while leaving genome sequences unchanged. It has been found to mediate gene regulation in relation to environmental conditions, neuronal activity and more (Bošković and Rando, 2018; Perez and Lehner, 2019). Germ cell-resident mechanisms that drive TEI have been found in both plants and animals, and include both nuclear (chromatin-based) as well as cytoplasmic mechanisms (Bošković and Rando, 2018; Castel and Martienssen, 2013; Perez and Lehner, 2019). A class of molecules that has been clearly linked to TEI is that of small RNAs, most notably short-interfering RNAs (siRNAs) and Piwi-interacting RNAs (piRNAs) (Bošković and Rando, 2018). These act as sequence-specific co-factors for Argonaute proteins, which in turn can impose gene-regulatory effects upon base-pairing of the small RNA to a target transcript. Such effects can range from target RNA destabilization to chromatin modification (Castel and Martienssen, 2013; Hutvagner and Simard, 2008; Peters and Meister, 2007).

In *C. elegans*, small RNAs with an established role in TEI are the 22G RNAs (de Albuquerque et al., 2015; Buckley et al., 2012; Mao et al., 2015; Phillips et al., 2015; Wan et al., 2018; Xu et al., 2018). These small RNAs are generated by RNA-dependent RNA Polymerases (RdRPs), in a process that also requires *Mutator* proteins, such as MUT-16, and are bound by a highly diversified, worm-specific Argonaute (WAGO) family (Gu et al., 2009; Phillips et al., 2012; Yigit et al., 2006; Zhang et al., 2011). 22G RNAs represent secondary siRNAs, as they are generated in response to target recognition by so-called primary Argonaute proteins, such as the *C. elegans* Piwi protein PRG-1, or the Argonaute protein that mediates exogenous RNAi: RDE-1 (Das et al., 2008; Yigit et al., 2006). However, following establishment, 22G RNA responses can be inherited across multiple generations, in some cases close to indefinitely, in absence of the original trigger (Luteijn and Ketting, 2013). Interestingly, both maternal as well as paternal inheritance of such responses has been described (Alcazar et al., 2008; Grishok et al., 2000; Lev et al., 2019; Wan et al., 2018). Finally, inheritance of 22G RNAs plays an important role in maintaining an endogenous 22G RNA pool that does not target essential genes. In absence of maternal and paternal 22G RNA inheritance severe sterility can develop, caused by 22G RNAs that erroneously silence genes required for germline development (de Albuquerque et al., 2015; Phillips et al., 2015). This phenotype is named *Mutator Induced Sterility* (Mis).

Molecular mechanisms linked to small RNA pathways are often organized in non-membranous organelles (Voronina et al., 2011), that often form through phase separation, driven by proteins with intrinsically disordered regions (IDRs), and/or proteins that are multivalent with regard to interactions with other proteins and RNA (Banani et al., 2017). In fact, the IDRs and multivalent character of these molecules are key characteristics for enhancing their phase separating behavior. In *C. elegans*, three such phase-separated condensates have been described to accommodate and affect small RNA-driven pathways: P granules, Z granules and *Mutator* foci (Phillips et al., 2012; Updike and Strome, 2010; Wan et al., 2018). Interestingly, none of these condensates has been found in mature sperm. In fact, it has been demonstrated that P granules dissociate during spermatogenesis (Updike and Strome, 2010). In contrast, P granules and Z granules have been closely connected to maternal TEI in *C. elegans* (Wan et al., 2018). In mammals, a dedicated paternal condensate has been described: the chromatoid body (CB) (Kotaja and Sassone-corsi, 2007). However, its function relates to RNA decay and translational repression during spermatogenesis, not to inheritance of specific factors. Finally, biomolecular condensates are well-described throughout germ cell development, but no such structures have been identified in mature sperm of any organism (Voronina et al., 2011). Possibly, this is linked to the fact that spermatogenesis is accompanied by a massive reduction of the intracellular content (Ellis and Stanfield, 2014). This also happens in *C. elegans*: during meiosis II a residual body is formed into which much of the spermatid cytoplasm is discarded, including, for instance, the spermatogenesis-expressed Argonaute protein ALG-3 (Conine et al., 2010).

Given the fact that much of the cytoplasmic content of spermatids is lost before spermatozoa are formed, one may question if paternal TEI can be mediated through the cytoplasm, or whether such inheritance via the father is restricted to chromatin-based mechanisms. Here we show that this is not the case. We identify the cytoplasmic Argonaute protein WAGO-3 as a factor required for paternal TEI and demonstrate that it is maintained within the cytoplasm of mature sperm. We identify the protein PEI-1, containing a BTB-BACK domain, as well as an IDR, as a factor that drives the formation of sperm-specific condensates (PEI granules), to which WAGO-3, but no other tested WAGO protein, is attracted. PEI granules are juxtaposed to membranous organelles such as mitochondria, and like mitochondria, depend on myosin-dependent transport to be maintained with the maturing sperm cell. This process critically depends on S-palmitoylation, suggesting that PEI granules are coupled to membranes by a lipid-anchor. We find that *pei-1*-like genes are also found in human, where they are expressed in, and have functions related to spermatogenesis, suggesting that the mechanism we here identify is broadly conserved.

## RESULTS

### WAGO-3 is required for an immortal germline and associates with paternal small RNAs

Many RNAi factors of *C. elegans* have been shown to be needed for proper germ cell development across generations, or germline immortality (Perez and Lehner, 2019). We found that WAGO-3 is also required for germline immortality, as out-crossed *wago-3* mutant animals became sterile in subsequent generations (Figure 1A), implicating WAGO-3 in a transgenerational mechanism that maintains germ cell health. We also analyzed how WAGO-3 affects the expression of a germline-specific mCherry::H2B transgene whose silencing can be initiated by the *C. elegans* Piwi protein PRG-1 (Bagijn et al., 2012; Lee et al., 2012; Ozata et al., 2019). Once established, this silencing can be maintained in absence of PRG-1 for many generations and is referred to as RNAe (RNAi-induced epigenetic silencing) (Ozata et al., 2019). We crossed an RNAe-silenced version of this transgene into *wago-3* mutants and analyzed its activity by microscopy. We found that WAGO-3 is not acutely required for RNAe, but that it is required for the stable transgenerational maintenance of the RNAe-driven silencing of this transgene (Figure 1B), indicating that WAGO-3 has a role in TEI.

**Figure 1.**
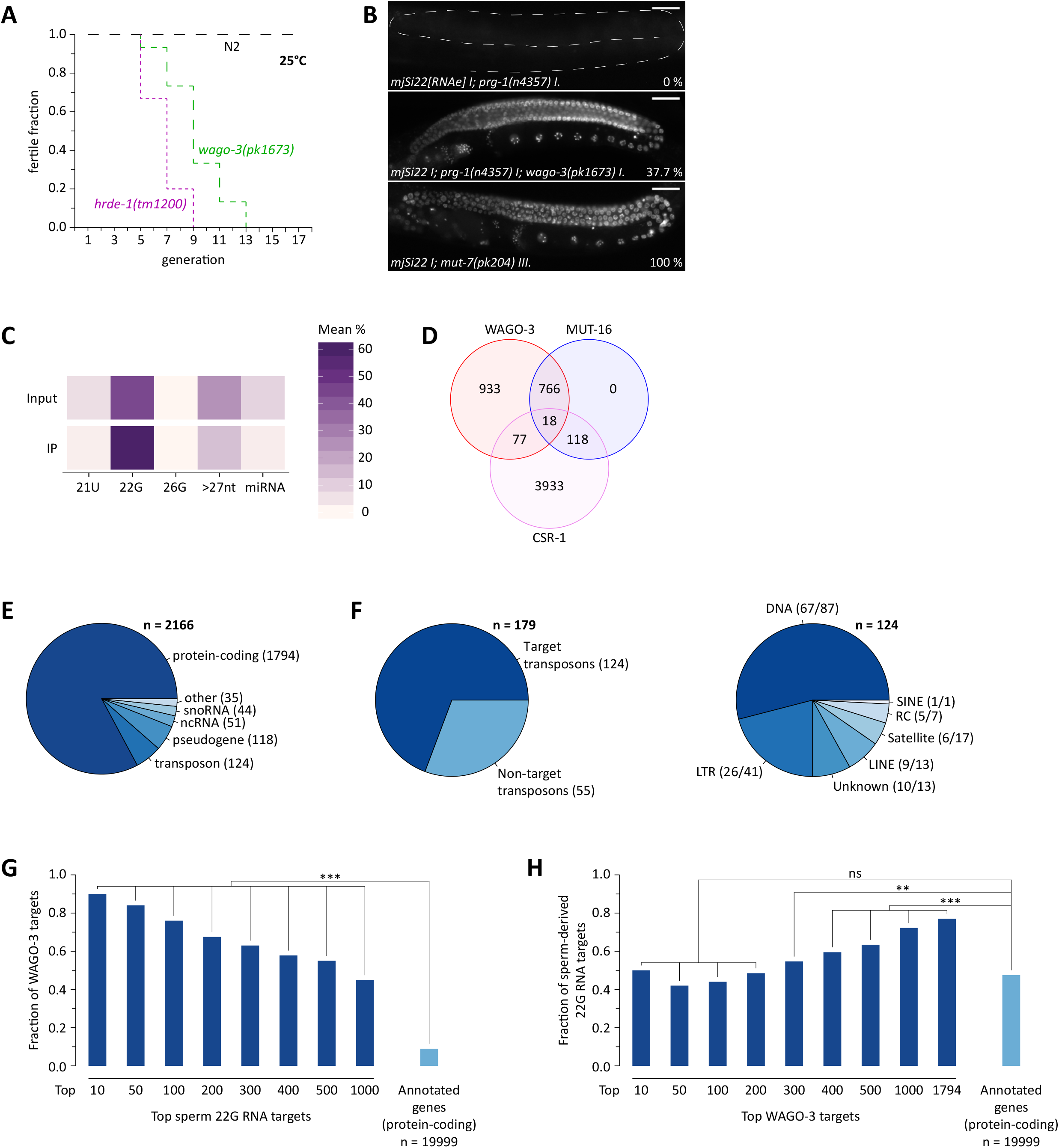
WAGO-3 is required for an immortal germline and associates with paternal small RNAs. A. Mortal germline assay representing loss of fertility of out-crosses strains with indicated genotype at 25°C. B. Fluorescence microscopy showing re-activation of a piRNA-targeted transgene in absence of both PRG-1 and WAGO-3. Expression of the transgene was determined after seven generations of homozygosity. No activity was detected in the first double mutant generation. Percentage in lower right corner shows fraction of animals expressing the transgene (n ≥ 50). Scale bars: 30 µm. C. Mean abundance of defined small RNA populations in the indicated small RNA libraries. D. Venn diagram showing overlap of protein-coding target genes between WAGO-3, CSR-1 and the *Mutator* complex. E. Distribution of WAGO-3 target genes. F. Genomic coverage and distribution of WAGO-3 targeted transposons. G. Bar chart showing how many of the top sperm-derived 22G RNA target genes (protein-coding), or of all annotated protein-coding genes, were also identified as WAGO-3 target genes. Statistical significance was tested with a Chi-square test (***: p ≤ 0.001). H. Bar chart showing how many of the top WAGO-3 target genes (protein-coding), or of all annotated protein-coding genes, were also identified as sperm-derived 22G RNA target genes. Statistical significance was tested with a Chi-square test (***: p ≤ 0.001, **: p ≤ 0.01, ns: p > 0.05). See also Figure S1.

Next, we aimed to determine the endogenous targets of WAGO-3, by sequencing WAGO-3-bound small RNAs. To achieve this, we inserted a *gfp::3xflag* encoding sequence directly downstream of the endogenous start codon of *wago-3*, to enable immunoprecipitation (IP) experiments. Following IP from adult animals, we sequenced small RNAs from both input and anti-FLAG IP samples. WAGO-3 is predominantly associated with small RNAs that have characteristics of 22G RNAs (Gu et al., 2009) (Figures 1C and S1A). Based on sequence complementarity we identified 2166 WAGO-3 targets. These targets cover the vast majority of known *Mutator* targets, while WAGO-3 and CSR-1 targets were almost mutually exclusive (Claycomb et al., 2009; Phillips et al., 2012) (Figure 1D). Notably, nearly 70 % of all annotated transposable elements were represented in WAGO-3-bound 22G RNAs (Figures 1E and 1F), consistent with its role in TE silencing (Robert et al., 2005; Vastenhouw et al., 2003). The vast majority of WAGO-3 associated 22G RNAs (83%) target exonic sequences over the whole gene body of both germline-expressed and soma-expressed protein-coding genes (Figures 1E and S1B-S1D). Among germline-expressed genes, WAGO-3 displayed a small, but significant preference for genes that are spermatogenic over genes that are oogenic (Figure S1B). Finally, we found a significant overlap of protein-coding target genes between WAGO-3 and sperm-derived 22G RNAs (Stoeckius et al., 2014) (Figure 1G and 1H). We note that these numbers are most likely underestimates since the sperm used to determine small RNA target genes was contaminated with 30 % spermatocytes (Stoeckius et al., 2014), and thus also included WAGO-1, CSR-1 and possibly other Argonaute associated 22G RNAs (Figures S2G-S2I). We conclude that WAGO-3 is involved in TEI and binds to 22G RNAs known to be paternally transmitted to the subsequent generation.

### WAGO-3 is expressed throughout germline development

We performed confocal microscopy to determine the expression pattern of GFP::3xFLAG::WAGO-3 (from here on referred to as WAGO-3). We detected specific and global expression throughout germline development, with localization to P granules in mitotic, meiotic and primordial germ cells (Figures S2A and S2B). Notably, we also found strong WAGO-3 signals within the spermatheca, where spermatozoa are stored, indicating that WAGO-3 is also expressed during spermatogenesis and that it is maintained in mature sperm.

### WAGO-3 is guided into sperm by PEI-1

Immunoprecipitation of WAGO-3 from late-L4 stage hermaphrodites, a stage during which spermatogenesis is ongoing, followed by label-free quantitative mass spectrometry (IP-MS/MS) identified a number of WAGO-3 co-enriched proteins. Among them, known P granule components like DEPS-1, PRG-1 and WAGO-1 were identified (Figure 2A). We focused, however, on the protein F27C8.5, which we named PEI-1 (<u>P</u>aternal <u>E</u>pigenetic <u>I</u>nheritance defective-1) (Figures 2A and 2B). We generated endogenously tagged *pei-1* alleles and found that PEI-1::mTagRFP-T (from here on referred to as PEI-1) was exclusively expressed during later stages of spermatogenesis, both in males (Figure S2C) as well as in L4 hermaphrodites (Figure S2D).

**Figure 2.**
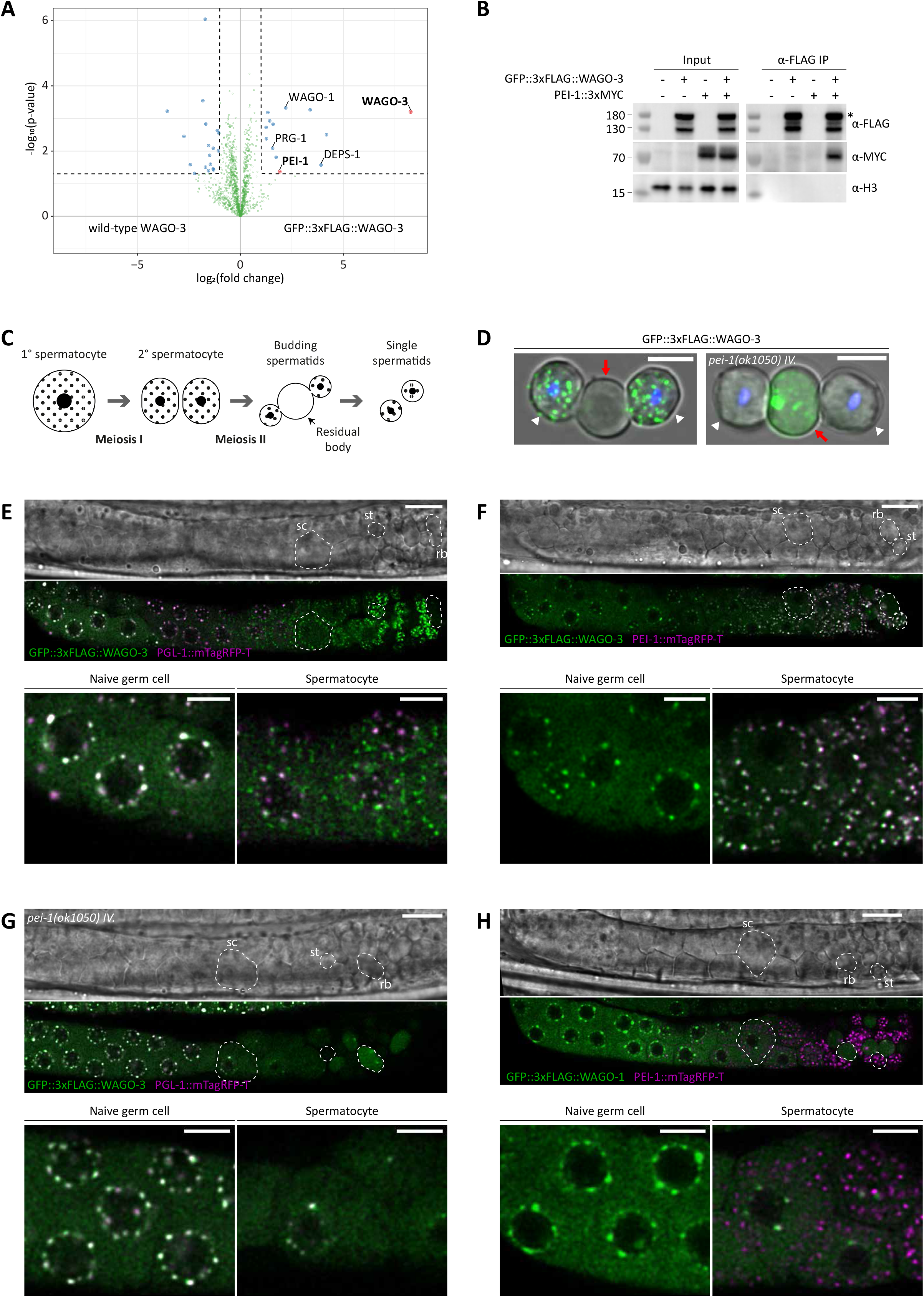
WAGO-3 is guided into sperm by PEI-1. A. Volcano plot representing label-free proteomic quantification of GFP::3xFLAG::WAGO-3 immunoprecipitation experiments from late-L4 stage hermaphrodite extracts. The *X*-axis indicates the mean fold enrichment of individual proteins in the control (wild-type WAGO-3) versus the genome-edited strain (GFP::3xFLAG::WAGO-3). The *Y*-axis represents −log_10_(p-value) of observed enrichments. Dashed lines show thresholds at P = 0.05 and twofold enrichment. Blue and green data points represent above and below threshold, respectively. WAGO-3 and PEI-1 are highlighted with red data points. B. Co-immunoprecipitation experiments using whole-worm extracts of late-L4 stage hermaphrodites. The asterisk marks the predicted full-length GFP::3xFLAG::WAGO-3 fusion-protein. C. Schematic summarizing spermatogenesis in *C. elegans*. D. Confocal maximum intensity projections of male-derived budding spermatids expressing GFP::3xFLAG::WAGO-3 in presence and absence of PEI-1. White arrow heads mark budding spermatids, red arrows indicate residual bodies. Hoechst33342 was used to stain DNA. Scale bars: 4 µm. E-H. Confocal micrographs showing spermatogenesis of late-L4 stage hermaphrodites expressing indicated proteins. sc – spermatocyte, rb – residual body, st – spermatid. Scale bars: 10 µm (proximal gonad), 4 µm (zoom). See also Figures S2 and S3.

We analyzed spermatogenic gonads of late-L4 stage hermaphrodites to look at PEI-1 and WAGO-3 expression more closely. In naïve germ cells, WAGO-3 localized to peri-nuclear P granules, marked by PGL-1 (Figure S2A). However, subcellular alterations occurred from the primary spermatocyte stage onwards. First, WAGO-3 was found to accumulate in distinct, non-peri-nuclear, cytoplasmic foci (Figures 2C-2E and S2E). Second, as previously described (Updike and Strome, 2010), P granules began to disappear (Figures 2E and S2E). Third, PEI-1 started to be expressed and always co-localized with WAGO-3 in cytoplasmic foci (Figures 2F and S2D). Using a publicly available deletion allele, we assessed the requirement of PEI-1 for the subcellular localization of WAGO-3 during sperm cell maturation. In *pei-1(ok1050)* mutants, WAGO-3 was absent from spermatozoa and instead was found in the residual body (Figures 2D; 2G; S2F), which contains discarded material that is not needed in mature sperm (Ellis and Stanfield, 2014). Thereby WAGO-3 followed the same fate as other Argonaute proteins like WAGO-1, ALG-3 and CSR-1 (Figures 2H and S2G-S2I). Conversely, loss of WAGO-3 did not affect PEI-1 localization (Figure S3). We conclude that WAGO-3 is special amongst *C. elegans* Argonaute proteins, as it is maintained in maturing sperm by re-localizing to cytoplasmic foci before P granules disappear. Furthermore, PEI-1 is a novel protein that marks these cytoplasmic foci and is required for proper WAGO-3 localization, and segregation into mature sperm.

### WAGO-3 and PEI-1 are crucial for paternal epigenetic inheritance of silencing information

We implemented a genetic system to test the roles of WAGO-3 and PEI-1 in parental TEI in an endogenous setting. We and others have previously shown that *C. elegans* individuals need parental 22G RNAs to specify the correct targets, particularly when also piRNAs are absent: Absence of parental 22G RNAs in animals that lack piRNAs but can make 22G RNAs results in sterility due to inappropriate gene silencing (Figure 3A). This phenotype was named *Mutator Induced Sterility* (Mis) (de Albuquerque et al., 2015; Phillips et al., 2015). We used this Mis phenotype to assess the ability of sperm or oocyte to provide silencing information that could lead to a rescue of the sterility. First, we tested whether either maternal or paternal 22G RNAs would be sufficient to prevent the Mis phenotype. As expected, this was indeed what we found (Figure 3B-C, top two bars). We next tested whether WAGO-3 or PEI-1 were required for either paternal and/or maternal rescue of the Mis phenotype. The offspring of both *wago-3*, as well as *pei-1* mutant males showed high degrees of sterility, similar to that observed from *mut-16* mutant males (Figure 3B), indicating that WAGO-3 and PEI-1 are required for paternal rescue of the Mis phenotype. In contrast, neither PEI-1 nor WAGO-3 was required for maternal rescue of this phenotype, whereas loss of *mut-7* did trigger sterility (Figure 3C). We conclude that PEI-1 and WAGO-3 are specifically required in the male, to provide embryos with paternal silencing memory that guides 22G RNA production in the offspring.

**Figure 3.**
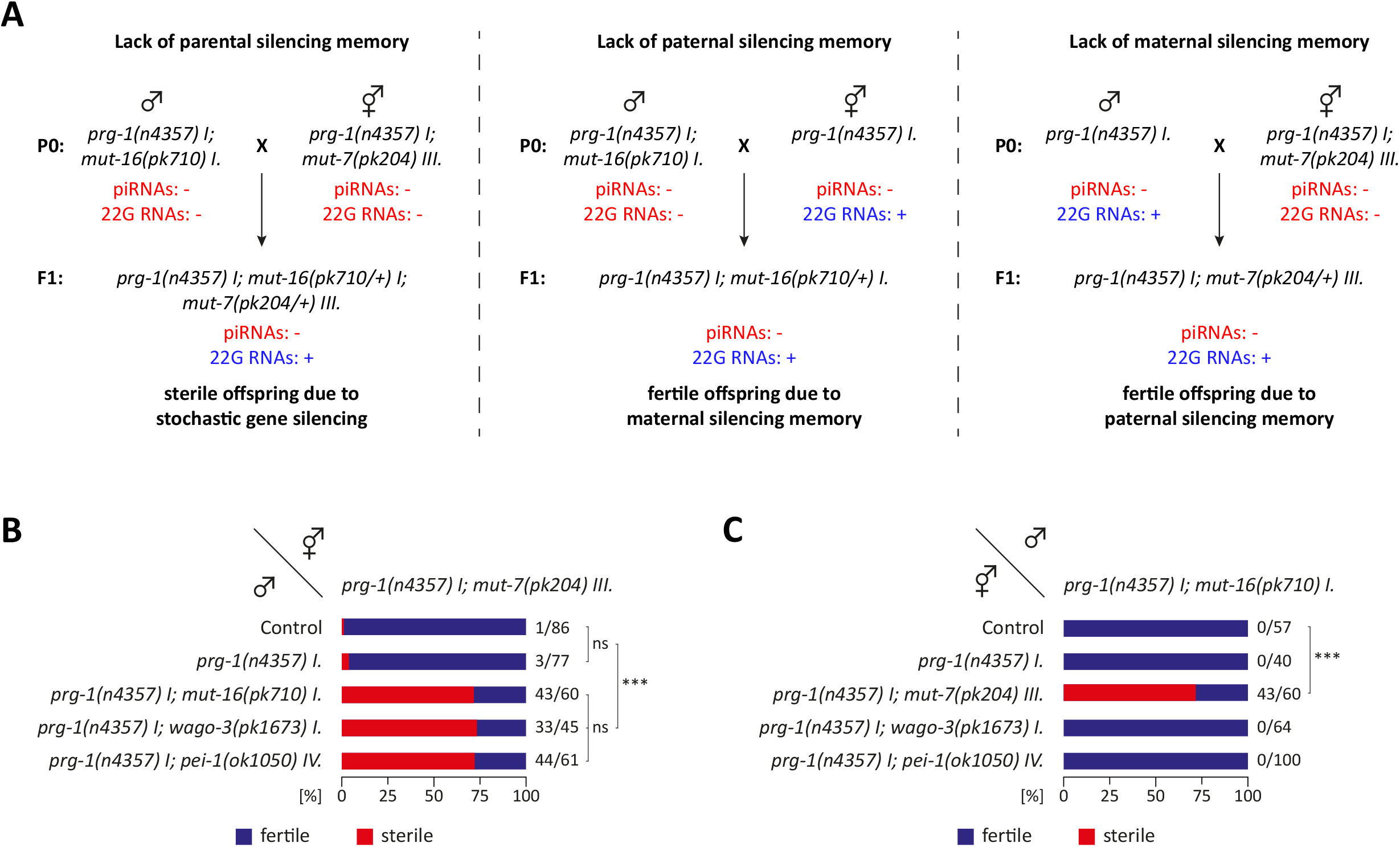
WAGO-3 and PEI-1 are crucial for paternal epigenetic inheritance of silencing information. A. Schematic illustrating the generation of animals that establish 22G RNA populations *de novo* in absence of piRNAs. Loss of MUT-7 or MUT-16 causes global depletion of WAGO-dependent 22G RNAs, whereas PRG-1 is required for piRNA biogenesis and function. B-C. Percentage of fertile F1 animals generated by indicated crosses. Fertility means: presence of maternal and/or paternal 22G RNA-mediated TEI. Sterility means: no 22G RNA-mediated TEI. Statistical significance was tested with a Chi-square test (***: p ≤ 0.001, ns: p > 0.05).

### Granule formation and WAGO-3 interaction are mediated via different PEI-1 domains

PEI-1 has a bimodal composition in terms of intrinsically ordered and intrinsically disordered regions. The N-terminal region is predicted to predominantly fold into alpha helical structures and likely adopts a BTB fold followed by a BACK domain (Figure 4A). While the BTB domain is described to mediate protein-protein interactions (Collins et al., 2001), the molecular function of the BACK domain remains speculative (Stogios and Privé, 2004). The C-terminal part of PEI-1 is predicted to be intrinsically disordered (Figure 4A). Following these predictions, we further edited the endogenous *pei-1::mTagRfp-t* locus by introducing defined deletions to generate six different, fluorescently tagged PEI-1 variants (Figure 4B). We then performed co-localization studies with WAGO-3 to determine effects on its subcellular distribution (Figure 4C). Deletion of the BTB domain resulted in normally appearing foci, which were accompanied by a diffuse cytoplasmic signal of both WAGO-3 and PEI-1 in spermatozoa. Loss of the BACK domain led to a similar phenotype, but in addition affected the number and homogeneity of cytoplasmic foci throughout spermatogenesis. PEI-1 lacking both the BTB and BACK domain caused a more severe phenotype. Although some foci were still present in maturing sperm, no foci, and only diffuse signals were detectable in spermatozoa. We note that none of these three deletion mutants resulted in PEI-1 or WAGO-3 accumulation in the residual body, and that both proteins always co-localized.

**Figure 4.**
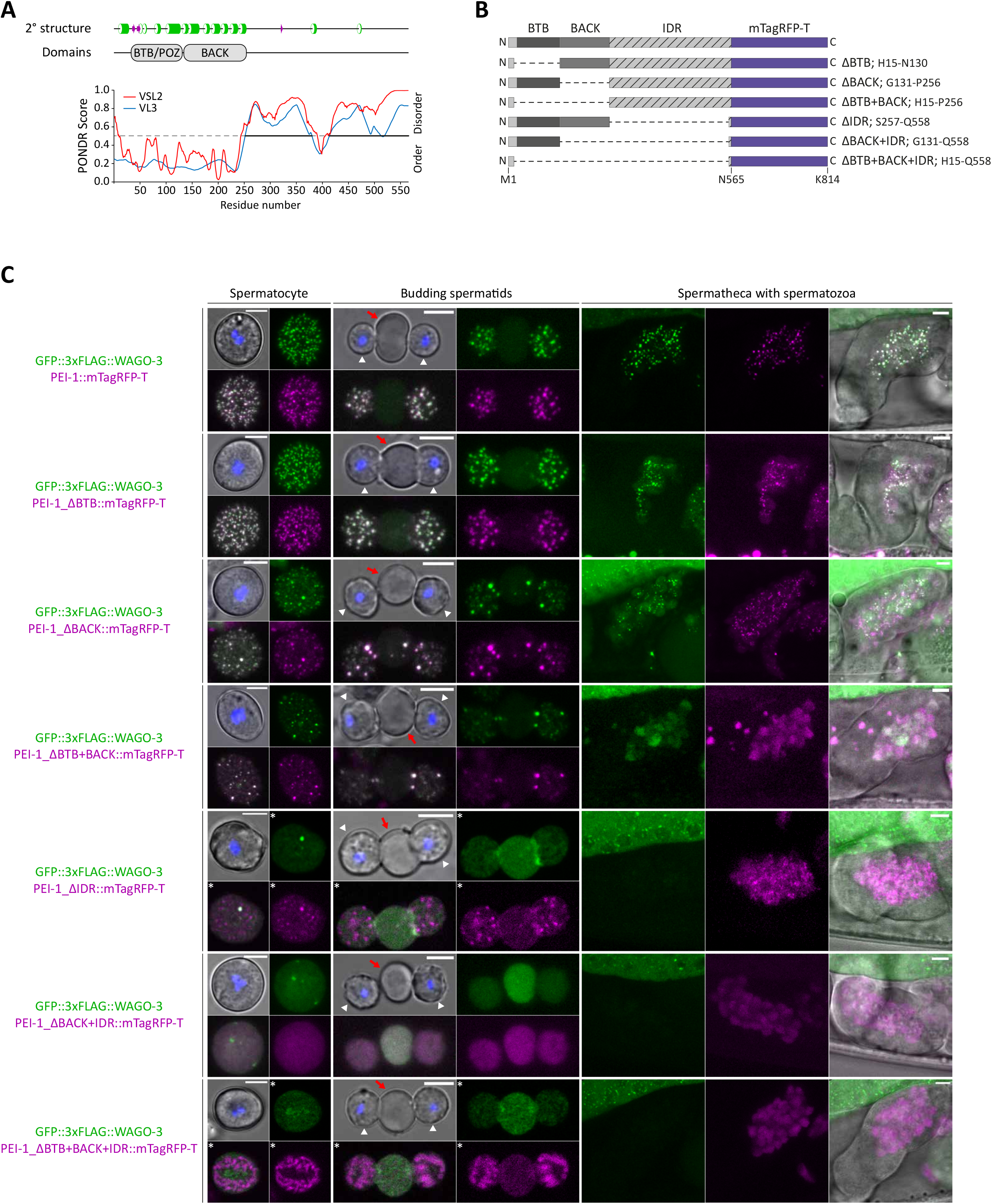
Granule formation and WAGO-3 interaction are mediated via different PEI-1 domains. A. PEI-1 protein composition in terms of secondary and fixed tertiary structure. Prediction of naturally disordered regions was performed using PONDR VSL2 and PONDR VL3 algorithms. Secondary structure elements were predicted using Jpred4 (Drozdetskiy et al., 2015). Helices are marked as green tubes, and sheets as magenta arrows. B. Schematic representation of truncated PEI-1::mTagRFP-T proteins generated by CRISPR/Cas9 mediated genome editing. IDR – intrinsically disordered region. C. Confocal maximum intensity projections of isolated spermatocytes (male-derived), budding spermatids (male-derived) and spermatozoa within the spermatheca (hermaphrodite) expressing indicated proteins. White arrow heads mark budding spermatids, red arrows indicate residual bodies. Hoechst33342 was used to stain DNA. Asterisks indicate optical sections. Scale bars: 4 µm. See also Figures S4.

Upon deletion of the IDR of PEI-1, WAGO-3 displayed diffuse distribution in spermatocytes, localized to residual bodies and was absent from spermatozoa. We note that the enrichment in the residual body was possibly based on active sorting, as a transcriptional GFP::3xFLAG reporter from the endogenous *wago-3* locus was found to be evenly distributed over the residual body and the budding spermatids, resulting in GFP positive spermatozoa (Figures S4A-S4C). Intriguingly, the remaining, structurally ordered part of PEI-1 did not accumulate in the residual body, but localized to faint foci in maturing sperm cells, indicating that the remaining PEI-1 protein can form larger assemblies with correct segregation. Additional removal of the BACK domain did not further affect the subcellular localization of WAGO-3: it remained enriched in the residual body. However, the remaining BTB::mTagRFP-T fusion-protein no longer formed foci, and was evenly distributed between residual body and budding spermatids, reminiscent of the symmetric segregation of free GFP::3xFLAG. To our surprise, the largest deletion of PEI-1, which only left a few N- and C-terminal amino acids, was found to localize to unknown structures that were segregated into spermatids (Figures 4C and S4D-S4E). Interestingly, co-staining of mitochondria revealed a mutually exclusive localization pattern (Figures S4F and S4G). We conclude that the IDR of PEI-1 is essential to bind WAGO-3 and contributes to foci formation, while the BTB and BACK domains play an important role in forming and stabilizing PEI-1 foci during the entire process of spermatogenesis, and continue to do so in spermatozoa. Additionally, both the BACK domain and the IDR are required for the asymmetric sorting of PEI-1 into the spermatids.

### PEI granules are independent condensates that retain WAGO-3 via hydrophobic interactions

Membrane-less compartments, like P granules and *Mutator* foci, are phase-separated condensates known to play crucial roles in germ cell biology (Phillips et al., 2012; Updike and Strome, 2010). We tested whether PEI-1 foci depend on these condensates by removing MUT-16, disrupting *Mutator* foci, or DEPS-1, disrupting P granule assembly (Phillips et al., 2012; Spike et al., 2008). Neither MUT-16 nor DEPS-1 was required for proper localization of PEI-1 during spermatogenesis, indicating that the formation of PEI-1 foci does not depend on these known, phase-separated condensates (Figure S3). Hence, PEI-1 foci represent an independent, novel entity, which we will from here on refer to as PEI granules.

Phase separation can involve various types of interaction including pi/pi, cation/pi, electrostatic and hydrophobic interactions (Hyman et al., 2014). Aliphatic compounds, such as 1,6-hexanediol, were shown to disrupt weak hydrophobic interactions, and hence affect phase separation driven by such interactions (Kroschwald et al., 2017). Thus, we monitored WAGO-3 and PEI-1 in isolated male-derived spermatocytes and budding spermatids in the presence of different concentrations of 1,6-hexanediol (Figure 5A). WAGO-3 was found to be highly sensitive, as no PEI granule localization was detected anymore in the presence of merely 1.25 % 1,6-hexanediol. This indicates a major role for hydrophobic interactions in WAGO-3 localization to PEI granules. Of note, WAGO-3 relocated preferentially to the residual body upon 1,6-hexanediol treatment, revealing an ongoing drive of these cells to relocate proteins into the residual body. In contrast, PEI-1 foci were found to be more resistant, especially in budding spermatids where even a 5 % 1,6-hexanediol treatment did not cause disassembly of PEI granules. These results suggest that the interaction between WAGO-3 and PEI granules is mostly hydrophobic, while PEI granules themselves significantly depend on other, or additional types of interactions, possibly mediated by the BTB and BACK domains.

**Figure 5.**
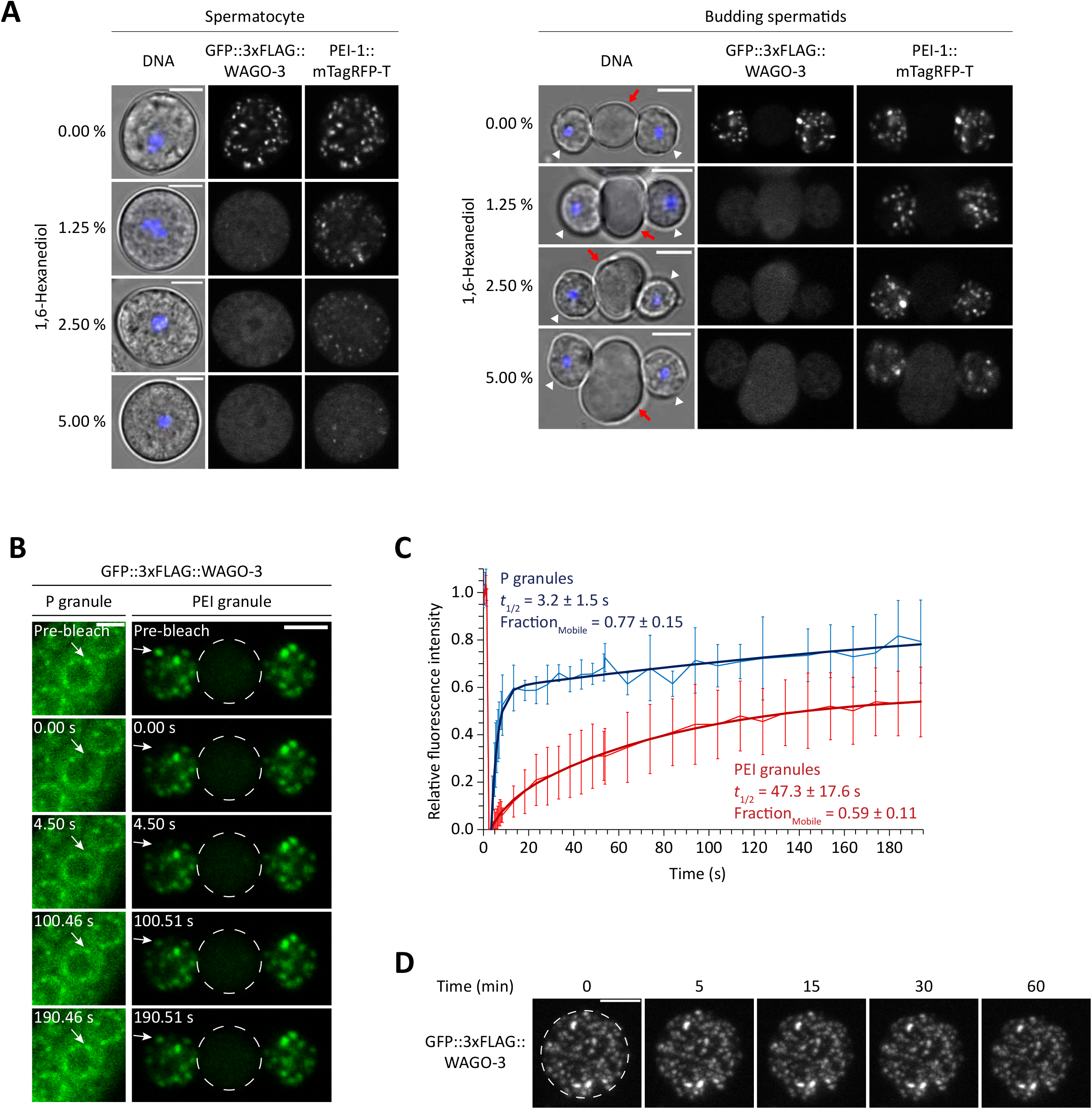
PEI granules are independent condensates that retain WAGO-3 via hydrophobic interactions. A. Confocal micrographs of isolated male-derived spermatocytes and budding spermatids after treatment with 1,6-hexanediol. White arrow heads mark budding spermatids, red arrows indicate residual bodies. Hoechst33342 was used to stain DNA. Scale bars: 4 µm. B. Time sequence showing fluorescence recovery after photobleaching (FRAP) of GFP::3xFLAG::WAGO-3 localizing to either P granules or PEI granules. Scale bars: 4 µm. C. FRAP recovery curve of GFP::3xFLAG::WAGO-3. Normalized data is presented as mean +/- SD and was fitted to a double exponential curve (n = 4 granules). D. Time sequence of GFP::3xFLAG::WAGO-3, taken from movie S1. Images are confocal maximum intensity projections of an isolated male-derived spermatocyte, which is marked by a dashed circle. Scale bar: 4 μm. See also Figure S3 and S5 and Movie S1.

To better understand the physical properties of PEI granules, we asked to which extent they exchange material with the cytoplasm. Therefore, we measured the fluorescence recovery after photobleaching (FRAP) of WAGO-3 in both P granules and PEI granules (Figures 5B and 5C). As previously reported, proteins localizing to the liquid-phase of P granules exhibit high recovery rates (Brangwynne et al., 2009; Putnam et al., 2019). Similarly, we found that WAGO-3 also showed very rapid fluorescence recovery in P granules (*t*_1/2_ = 3.2 +/- 1.5 s). Strikingly, WAGO-3 exhibited much slower exchange dynamics when localizing to PEI granules in budding spermatids (*t*_1/2_ = 47.3 +/- 17.6 s). Moreover, we found that the mobile fraction of WAGO-3 within the condensates is slightly reduced in PEI granules compared to P granules. This suggests that PEI granules are more gel-like than P-granules. The prevalence of certain amino acids has been shown to modulate the material property of phase-separated condensates (Wang et al., 2018). In particular, glycine residues were shown to maintain liquidity, while serine and glutamine residues promote hardening of condensates. Thus, we analyzed the amino acid composition of the intrinsically disordered regions that were predicted for PEI-1, and compared that to IDR compositions of PGL-1 and PGL-3, both known to localize to the liquid phase of P granules (Hanazawa et al., 2011; Kawasaki et al., 1998, 2004), and of MEG-3 and MEG-4, both reported to form gel-like assemblies (Putnam et al., 2019) (Figure S5). While glutamine was not found to be strongly enriched in any of the IDRs, serine was significantly enriched in all of them, and most highly in PEI-1, MEG-3 and MEG-4. In addition, PGL-1 and PGL-3 IDRs were strongly enriched for glycine, whereas such enrichment was absent from PEI-1, MEG-3 and MEG-4 IDRs. This similar amino acid profile between PEI-1 and MEG-3/4 IDRs is consistent with the idea that PEI granules are more gel-like than P-granules.

Finally, we live-imaged PEI granules by monitoring WAGO-3, and found that they were rather static. We did not observe any major movements in a period of one hour (Figure 5D and Movie S1), suggesting an attachment to bigger structures, which prevents any fusion or fission events between individual foci. Everything considered, we conclude that PEI granules are stable, independent condensates that retain WAGO-3 via hydrophobic interactions.

### Segregation of PEI granules depends on membranous organelle transport

We next addressed the question of how PEI granules are transported into budding spermatids. A substantial amount of cellular material, including free ribosomes, the endoplasmic reticulum and the Golgi apparatus, is asymmetrically segregated into residual bodies during the second meiotic division. Only a few organelles, like the nucleus, mitochondria and the sperm-specific fibrous body-membranous organelles (FB-MOs) are exclusively sorted into budding spermatids (Ellis and Stanfield, 2014). Given the very low mobility of PEI granules, we reasoned that they might be associated with one of these membranous organelles. A previous study showed that the myosin VI motor protein SPE-15 is required for proper segregation of mitochondria and FB-MOs into budding spermatids (Kelleher et al., 2000). Thus, we asked whether the asymmetric sorting of PEI granules is subject to the same principle. In absence of SPE-15, PEI granules were detected in both budding spermatids and residual bodies (Figure 6A), revealing that their subcellular distribution indeed depends on SPE-15, and is thus likely driven by the segregation of mitochondria and/or FB-MOs. We also performed confocal microscopy to determine the subcellular localization of mitochondria in maturing sperm cells in relation to PEI granules. This revealed PEI-1 localization close to mitochondria (Figure S6A; Movies S2 and S3), without showing any overlaps. To probe FB-MO association, we looked at WAGO-3 and SPE-45::mCherry localization in spermatids that budded off the residual body (Figure S6B). SPE-45 is a protein known to localize to the membranous organelles of FB-MOs, which are found close to the cell membrane in spermatids. While SPE-45::mCherry showed this peripheral distribution, WAGO-3 foci were detected throughout the cytoplasm, suggesting that WAGO-3 is not stably associated with FB-MOs.

**Figure 6.**
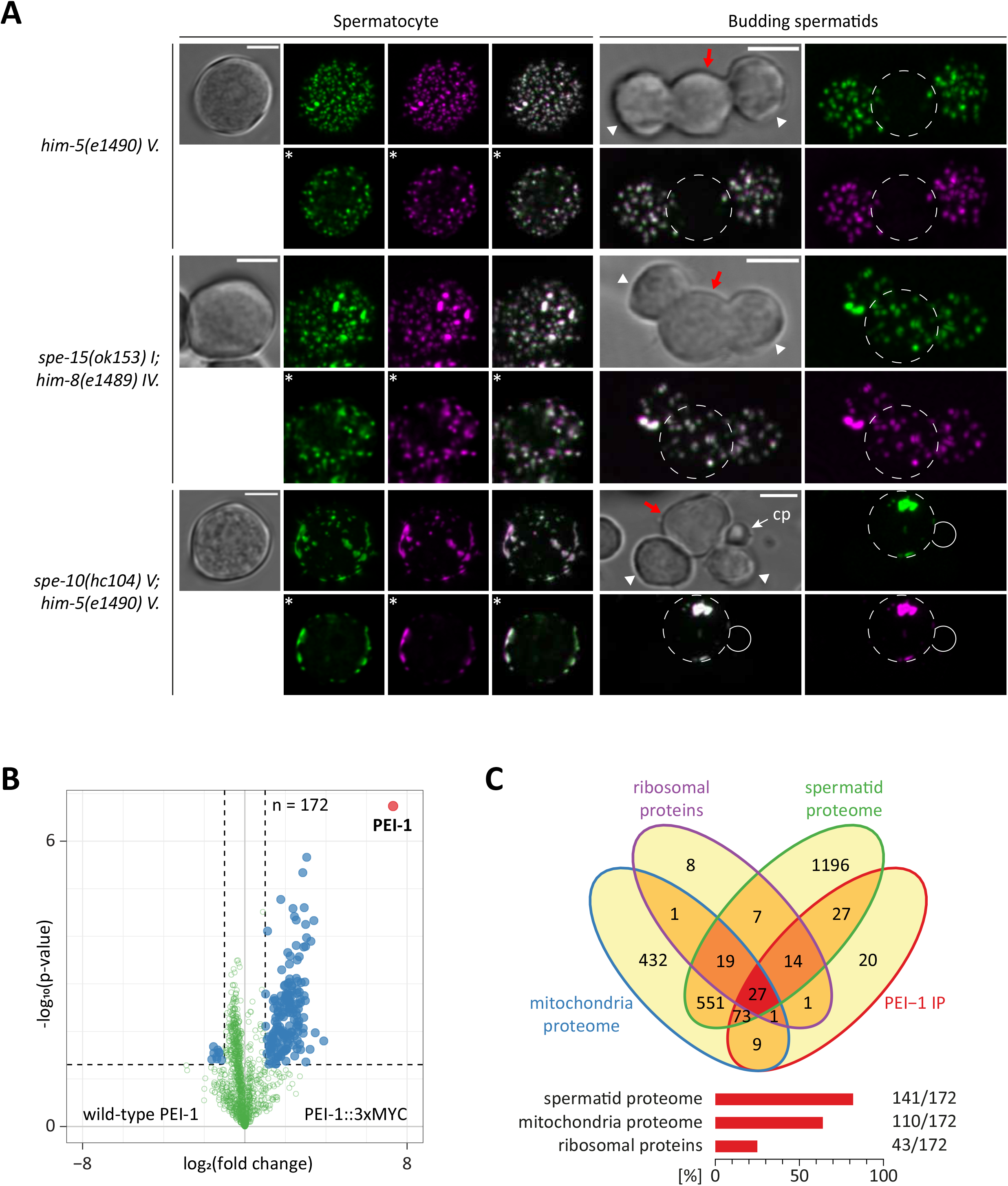
Segregation of PEI granules depends on organelle transport and S-palmitoylation. A. Confocal maximum intensity projections of isolated male-derived spermatocytes and budding spermatids expressing both GFP::3xFLAG::WAGO-3 (green) and PEI-1::mTagRFP-T (magenta) in indicated mutants. White arrow heads mark budding spermatids, red arrows and dashed circles indicate residual bodies. Asterisks indicate optical sections. cp – cytoplast. Scale bars: 4 µm. B. Volcano plot representing label-free proteomic quantification of PEI-1::3xMYC immunoprecipitation experiments from late-L4 stage hermaphrodite extracts. The *X*-axis indicates the mean fold enrichment of individual proteins in the control (wild-type PEI-1) versus the genome-edited strain (PEI-1::3xMYC). The *Y*-axis represents −log_10_(p-value) of observed enrichments. Dashed lines show thresholds at P = 0.05 and twofold enrichment. Blue and green data points represent above and below threshold, respectively. PEI-1 is highlighted with a red data point. C. Venn diagram and bar chart showing comparison of enriched proteins from (B) with published spermatid proteome, mitochondria proteome and ribosomal proteins. See also Figure S6 and Movies S2-S4.

Additional evidence for association of PEI granules with mitochondria comes from an IP-MS/MS experiment on late-L4 stage animals expressing PEI-1::3xMYC from the endogenous *pei-1* locus (Figures 6B and 6C). This experiment identified 172 enriched proteins. The vast majority (82 %), including PEI-1 and WAGO-3, were also detected in a proteomic analysis of male-derived spermatids (Ma et al., 2014). Strikingly, 110 proteins (64 %) were previously identified in a proteomic study on purified mitochondria (Jing et al., 2009). We also identified 43 ribosomal proteins (25 %). Both sets are fully consistent with interaction of PEI granules with mitochondria, as previous proteomic and microscopic analyses identified and visualized cytosolic ribosomes on the surface of mitochondria (Gold et al., 2017; Jing et al., 2009). We also identified major sperm protein (MSP), which is the main component of the fibrous body (FB) (Ellis and Stanfield, 2014). We note that MSP was also identified on isolated mitochondria (Jing et al., 2009), suggesting a common contamination or possible interaction of both organelles. Taken together, we propose that PEI granules interact with mitochondria, and possibly with FB-MOs, which transport PEI granules into spermatids in a myosin VI-dependent manner.

### Correct segregation of PEI granules requires S-palmitoylation

S-palmitoylation is a covalent attachment of palmitic acid to a protein and can serve as membrane anchor, guiding proteins to membranous structures. S-palmitoylation typically occurs on the Golgi complex or on Golgi-derived membranes (Tabaczar et al., 2017). SPE-10, a sperm-specific palmitoyltransferase, has been reported to localize to FB-MOs, and to be required for the stable interaction between the FB and the Golgi-derived membranous part (Gleason et al., 2006). In absence of SPE-10, defects of FB-MOs become apparent during the second meiotic division: the FBs dissociate prematurely from the membranous parts and end up in the residual body, where they frequently bud off in so-called cytoplasts. We found that PEI granules were severely defective in *spe-10* mutants. Already in spermatocytes, a stage well before the second meiotic division, PEI-1 localized to large and irregularly shaped patches, which were found arranged along the cell membrane (Figure 6A). These patches, like the much smaller wild-type PEI granules, were very static and did not show signs of fusion or fission (Movie S4). At later spermatogenic stages, large PEI-1 aggregates were detected in the residual body, leaving no detectable PEI-1 signal in the spermatids (Figure 6A). No PEI-1 signal was detected in cytoplasts (Figure 6A), suggesting that the PEI-1 aggregates in the residual body of *spe-10* mutants were not FB-associated. Our results show that the molecular function of SPE-10, i.e. S-palmitoylation, is not restricted to FB-MO stabilization, but also affects both shape and segregation of PEI granules. WAGO-3 and PEI-1 still co-localized in absence of SPE-10, indicating that S-palmitoylation does not affect the interaction between both proteins. Rather, we hypothesize that it mediates the association of PEI granules with mitochondria.

## DISCUSSION

Our work identifies WAGO-3 as a cytoplasmic Argonaute protein that is inherited via the sperm, and how WAGO-3 is secured within the maturing sperm cells by joining a sperm-specific condensate made by PEI-1. Our findings are summarized in Figure 7, but a number of additional aspects will be discussed below.

**Figure 7.**
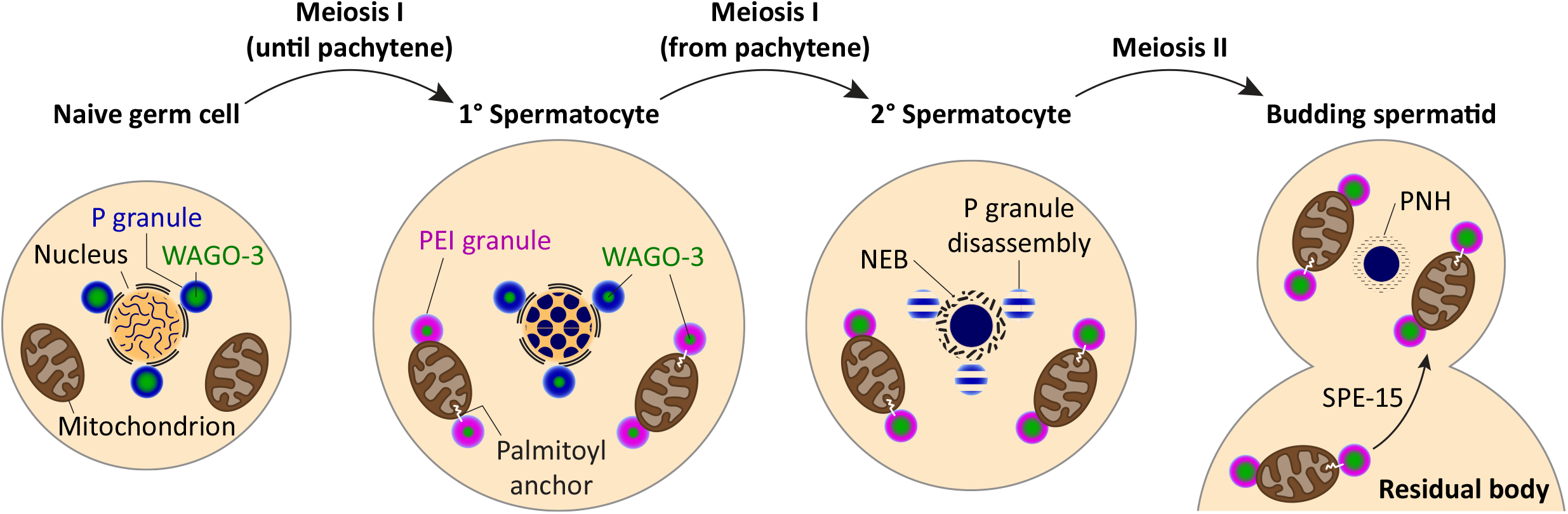
Model of paternal inheritance of WAGO-3. NEB – nuclear envelope breakdown, PNH – perinuclear halo

### PEI granule formation and WAGO specificity

We identified PEI-1, a conserved protein in terms of domain composition that is required for WAGO-3 localization in mature sperm. PEI-1 exclusively marks a novel, sperm-specific compartment, the PEI granule, which ensures proper subcellular segregation of WAGO-3. Both the IDR and the structurally ordered region of PEI-1 are able to form assemblies, and likely contribute to granule formation. However, we also found differences. The BTB and BACK domains of PEI-1 have a strong effect on the stability of PEI granules, whereas the IDR is essential for WAGO-3 recruitment. We note that BTB domains can mediate both homo- and heteromeric oligomerization (Collins et al., 2001), which provides multivalency, a property known to drive phase separation (Banani et al., 2017). We propose that the BTB and BACK domains stabilize the PEI-1 IDR-IDR interactions, allowing the formation of stable granules, whereas the IDR also specifies interactions with other proteins, such as WAGO-3.

Although a number of Argonaute proteins have been reported in spermatocytes (Batista et al., 2008; Buckley et al., 2012; Conine et al., 2010; Wan et al., 2018), we did not find any of these in spermatozoa within the spermatheca. WAGO-1 and CSR-1 have specifically been proposed to be maintained during sperm cell maturation (Conine et al., 2010, 2013), however, using tagged proteins expressed from endogenous loci, we could not confirm their presence in mature sperm. Especially WAGO-1 has a very similar protein sequence compared to WAGO-3, making the difference between WAGO-3 and WAGO-1 localization during sperm development remarkable. Further experiments will be required to identify the origin of this differential behavior.

### Transport of PEI granules

PEI granules are segregated to sperm via myosin VI dependent transport that is also known to be required for proper localization of mitochondria and FB-MOs (Kelleher et al., 2000). Our data suggest that mitochondria are an important vector of transport for the PEI granules, even though we cannot exclude a function for FB-MOs in this process. Interestingly, association of RNAi-related pathways with mitochondria has been described in various animals and include spermatogenic structures like pi-bodies, piP-bodies, chromatoid bodies and mitochondria-associated ER membranes (Wang et al., 2020). It appears that interaction with mitochondria may be used for various purposes in the context of small RNA pathways, ranging from small RNA biogenesis (Ge et al., 2019; Munafò et al., 2019) to the here identified role in transport and TEI.

An important question to further resolve is how PEI granules can interact with membranous structures. We show that SPE-10 mediated S-palmitoylation plays an important role in PEI granule localization, suggesting that anchoring of PEI granules may proceed via a lipid modification. It remains unknown whether PEI-1 itself is targeted for S-palmitoylation, but the striking subcellular localization of a large PEI-1 deletion that only maintains a few amino acids at both N- and C-termini provides an intriguing clue. The subcellular structures identified by this variant are not mitochondria. By principle of exclusion, they might well be FB-MOs. The observed structures indeed resemble FB-MO morphology, as revealed by electron microscopy (Fabig et al., 2019). This would imply that the most terminal amino acids of PEI contain information for FB-MO localization, where PEI-1, or an associated protein, could be palmitoylated by SPE-10 (Gleason et al., 2006). In this scenario, the large PEI-1 deletion visualizes a normally transient association of PEI-1 with FB-MOs, in order to interact with the SPE-10 enzyme. Interestingly, both the BACK domain and the IDR, which we show are required for proper spermatid localization of PEI-1, contain predicted palmitoylation sites (Figure S6C), suggesting that PEI-1 itself may be modified by SPE-10. More experiments will be needed to test these hypotheses.

### Why are PEI granules needed and how do they release their cargo in the embryo?

Why are PEI granules required for paternal TEI and how can they release their contents upon fertilization? The answer to the first question could be related to the fact that in most nematodes the nuclear membrane breaks down during late stages of spermatogenesis (Yushin and Malakhov, 2014). In mature sperm, the highly condensed genome is encapsulated by an RNA-protein halo (Ward et al., 1981). Since P granules are closely associated with the nuclear envelope, they may not be able to act as carrier of epigenetic information in sperm. Indeed, except WAGO-3, all described Argonaute proteins that were reported in P granules are excluded from mature sperm. Hence, a different, non nuclear envelope-associated condensate may be required to stabilize and keep specific proteins in maturing sperm. A different condensate may allow add specificity towards Argonaute inheritance. Whereas P granules are known to house many Argonaute proteins, only one, or a selected set may be appropriate to load into mature sperm. The reduced exchange dynamics of PEI granules compared to P granules might provide another important aspect for why PEI granules are required: PEI granules may be more viscous than P granules, allowing more robust transport. We do note that WAGO-3 recovery in PEI granules is still faster than that typically found in gel-like assemblies (Putnam et al., 2019). This may be related to the fact that in order to function in TEI, WAGO-3, together with its 22G RNA cofactors, has to be released into the oocyte, as paternal mitochondria are degraded during early development (Sato and Sato, 2017). We can envisage a number of possibilities of how this release may work. First, upon fertilization paternally inherited factors are massively diluted, which may cause PEI granule disassembly as concentrations drops below critical thresholds for phase separation. Thus, WAGO-3 proteins might efficiently diffuse into the zygote. Second, maternally deposited factors may stimulate WAGO-3 release by mediating post-translational protein modifications, which have been shown to regulate phase separation and affect physical properties of biomolecular condensates (Hofweber and Dormann, 2019). Third, S-palmitoylation is reversible (Tabaczar et al., 2017), raising the possibility that PEI granules as a whole are released after fertilization. It will be interesting to experimentally test these, and other possibilities for how paternal WAGO-3 can be used to prime embryonic 22G RNA biogenesis.

### PEI-1 related proteins in other species

How conserved is the mechanism we uncovered? Based on primary sequence, PEI-1 is nematode specific (Figure S7A). However, when domain organization is considered, PEI-1-related proteins can be easily identified, for example in human (Figure S7B). Particularly the human protein BTBD7, which is expressed in various tissues including testis, has a domain organization that closely resembles that of PEI-1. Interestingly BTBD7 carries a predicted myristoylation site close to its N-terminus, suggesting it may be membrane bound. The other human proteins shown in Figure S7B are also known to be expressed in testis, and for two of these, BTBD18 and GMCL1 (which is the human homolog of Drosophila gcl), functions during spermatogenesis have been described (Kleiman et al., 2003; Zhou et al., 2017). Interestingly, BTBD18 forms nuclear foci (Zhou et al., 2017), and GMCL1 has been described to interact with IDRs found in primate-specific GAGE proteins, and to affect their localization (Gjerstorff et al., 2012), similar to what we find for PEI-1 and WAGO-3. Hence, it seems likely that the mechanism we here reveal is broadly conserved in germ cell biology, and possibly also beyond.

## ACKNOWLEDGEMENTS

We thank all of the current and former members of the Ketting laboratory for helpful discussions and feedback on the manuscript. We are grateful to Helge Grosshans for critical reading of the manuscript. A special thanks to Miroslav Dörr and Svenja Hellmann for excellent technical and experimental support. Clara Werner of the Institute for Molecular Biology Genomics Core Facility is thanked for small RNA library preparation. We would like to thank the Institute for Molecular Biology Media Laboratory, Microscopy, Proteomic and Genomic Core Facilities for consumables, equipment and experimental support. Some strains were provided by the *Caenorhabditis* Genetics Center (CGC), which is funded by NIH Office of Research Infrastructure Programs (P40 OD010440). This work was supported by grants of the Deutsche Forschungsgemeinschaft KE 1888/1-1, KE1888/1-2 and KE 1888/6-1 (R.F.K.) and the National Institute of Health R35 GM119656 (C.M.P.), and T32 GM118289 (D.H.N.).

## AUTHOR CONTRIBUTIONS

J.S. and R.F.K. conceived the study and designed experiments. J.S. executed experiments and performed data analysis. S.D. and F.B. performed MS analysis. A.M.d.J.D. and A.S. performed smRNA-seq analysis. D.H.N. and C.M.P. provided USC988, USC1092 and USC1137 strains. E.J.G., H.L. and S.W.L. provided the SL1697 strain. R.F.K. supervised the project. J.S. and R.F.K. wrote the manuscript with input from all authors.

## METHODS

### *C. elegans* culture and strains

Unless otherwise stated, all worm strains were cultured according to standard laboratory conditions at 20°C on Nematode Growth Medium (NGM) plates seeded with *Escherichia coli* OP50 (Brenner, 1974). Animals for IP-MS/MS experiments were grown on egg plates (90 mm diameter) (Schweinsberg and Grant, 2013) for one generation, synchronized by bleaching, and then grown on standard NGM plates (90 mm diameter) for one generation before harvest. Egg plates were generated by thoroughly mixing egg yolk with 50 ml LB media/egg. Following incubation at 65°C for 2-3 hours, the mixture was allowed to cool to room temperature before adding 10 ml OP50 culture/egg. About 10 ml was put on top of standard NGM plates (90 mm diameter) and incubated at room temperature. Next day, excess liquid was decanted and egg plates were incubated at room temperature for another two days. All strains are in the N2 Bristol background. Strains used in this study are listed below.

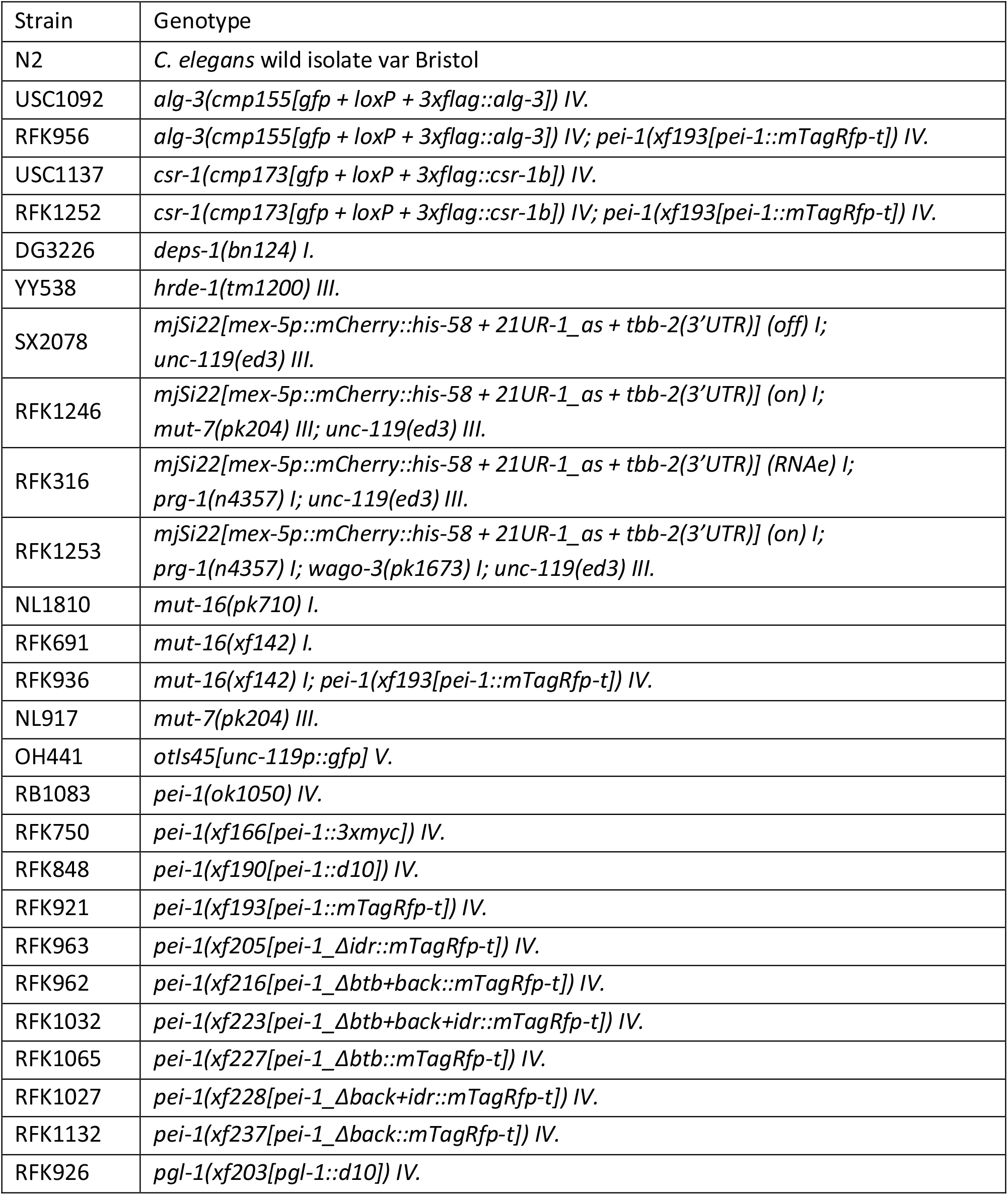
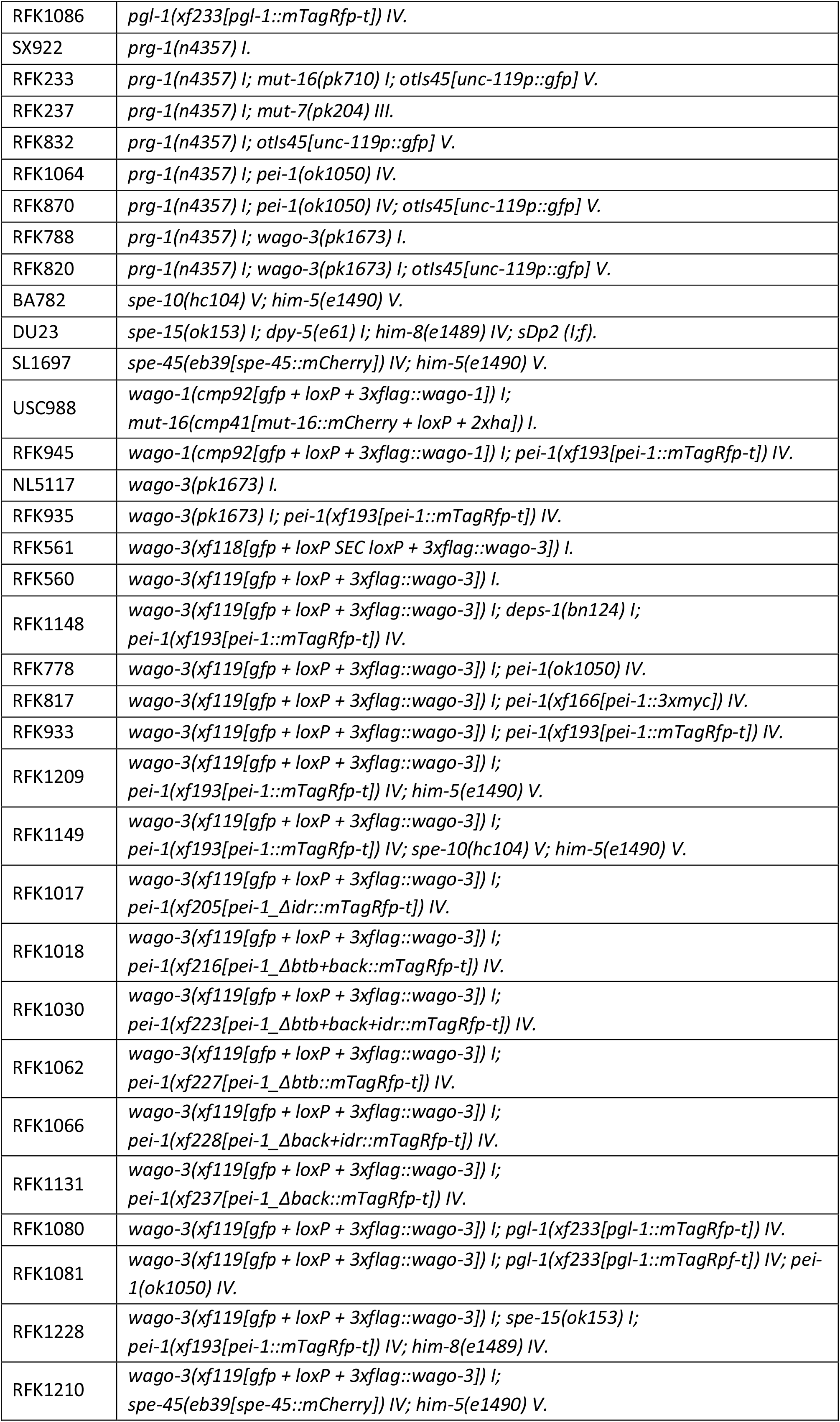
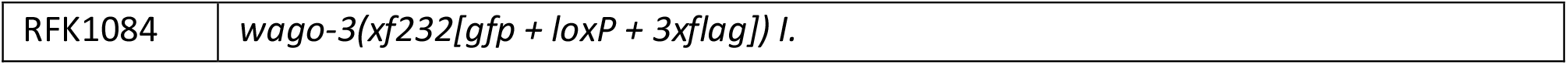

### Mortal germline assay

All mutant strains were confirmed and out-crossed four times before starting the experiment. For each strain, 90 L2 or L3 animals were distributed to 15 NGM plates (90 mm diameter), resulting in six larvae per plate. Animals were grown at 25°C. Worms were picked onto fresh plates every second generation. The experiment was stopped after 17 generations.

### *Mutator*-induced sterility crosses

All strains were confirmed and out-crossed two times before setting up crosses. We note that out-crossing ensured comparable results as an enhanced Mis phenotype was observed when using non-out-crossed animals. The transgenic allele *otIs45[unc-119p::gfp] V*. was always present in paternal strains and served as mating control to avoid picking self-fertilized offspring. Only L2 stage F1 animals were picked onto individual plates to avoid any biased selection. After three days, male or dead F1 animals were excluded from the analysis. Fertility of F1 animals was determined by the presence of F2 animals after another two to four days.

### CRISPR/Cas9-mediated genome editing

All protospacer sequences were chosen using CRISPOR (http://crispor.tefor.net) (Haeussler et al., 2016) and, unless otherwise stated, cloned in either pRK2411 (plasmid expressing Cas9 + sgRNA(F+E) (Chen et al., 2013); derived from pDD162) or pRK2412 (plasmid expressing sgRNA(F+E) (Chen et al., 2013) with Cas9 deleted; derived from pRK2411) via site-directed, ligase-independent mutagenesis (SLIM) (Chiu et al., 2004, 2008). pDD162 (Peft-3::Cas9 + Empty sgRNA) was a gift from Bob Goldstein (Addgene plasmid # 47549; http://n2t.net/addgene:47549; RRID:Addgene_47549) (Dickinson et al., 2013). SLIM reactions were transformed in Subcloning Efficiency™ DH5α™ Competent Cells (Art. No. 18265017, Invitrogen™) and plated on LB agar plates supplemented with 100 µg/ml ampicillin. All protospacer sequences are listed below.

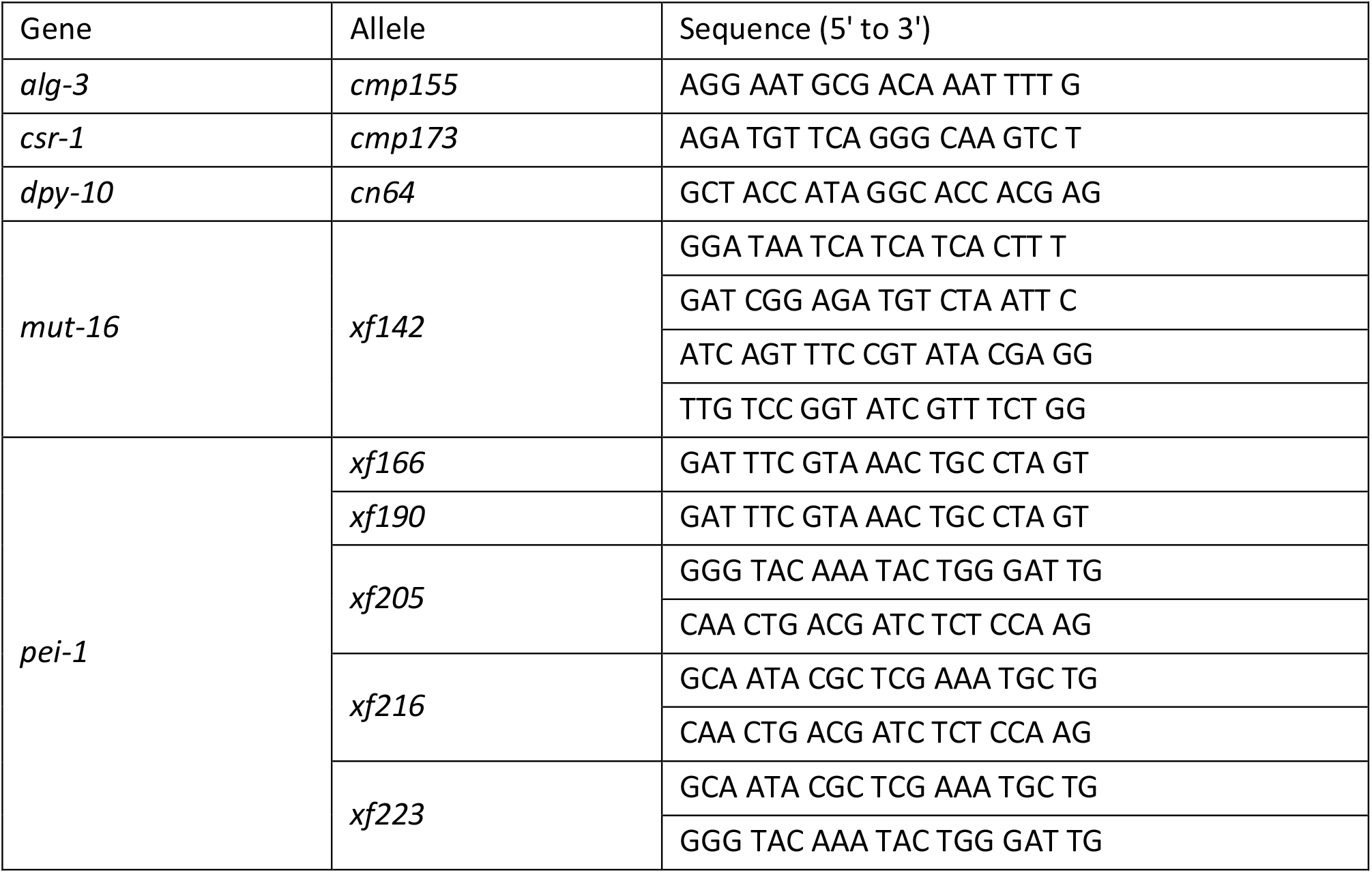
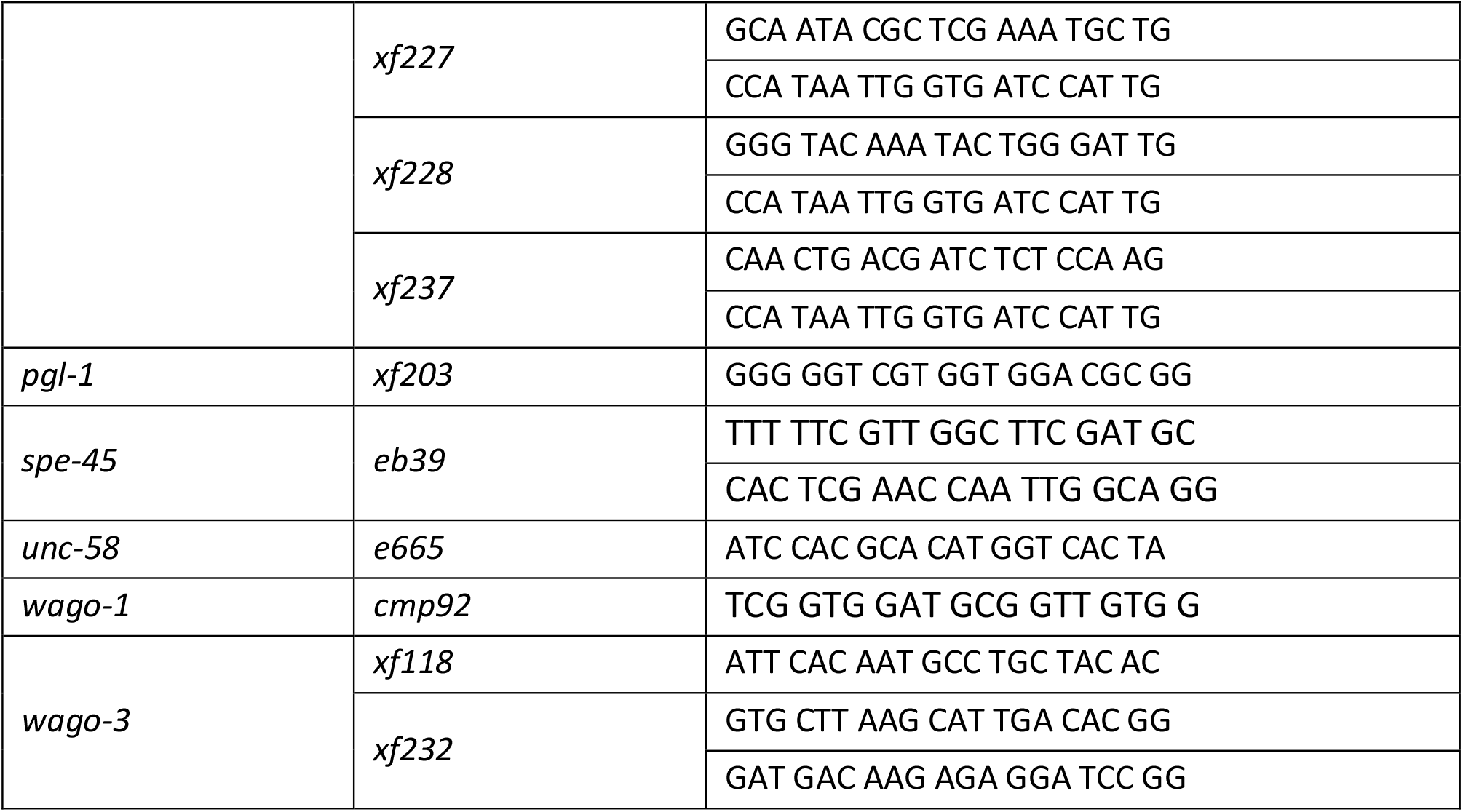

Insertions of a *gfp::3flag* sequence were based on plasmid DNA donor templates containing a self-excising drug selection casseste (SEC), which were designed and cloned as previously described (Dickinson et al., 2015). pDD282 was a gift from Bob Goldstein (Addgene plasmid # 66823; http://n2t.net/addgene:66823; RRID:Addgene_66823) (Dickinson et al., 2015). pJW1259 was used as Cas9 plasmid and was a gift from Jordan Ward (Addgene plasmid # 61251; http://n2t.net/addgene:61251; RRID:Addgene_61251) (Ward, 2014). pGH8, pCFJ90 and pCFJ104 served as co-injection markers and were gifts from Erik Jorgensen (Addgene plasmid # 19359; http://n2t.net/addgene:19359; RRID:Addgene_19359, Addgene plasmid # 19327; http://n2t.net/addgene:19327; RRID:Addgene_19327, Addgene plasmid # 19328; http://n2t.net/addgene:19328; RRID:Addgene_19328) (Frøkjær-Jensen et al., 2008). All plasmids were purified from 4 ml bacterial culture using either NucleoSpin^®^ Plasmid (REF 740588.50, Macherey-Nagel^®^) or PureLink™ HiPure Plasmid Miniprep Kit (Art. No. K210011, Invitrogen™), eluted in sterile water and confirmed by enzymatic digestion and sequencing.

PCR products served as linear, double-stranded DNA donor templates for the insertion of *mTagRfp-t* and *mCherry* sequences. The *mTagRfp-t* coding sequence including three introns and flanking homology regions was amplified from pDD286, which was a gift from Bob Goldstein (Addgene plasmid # 70684; http://n2t.net/addgene:70684; RRID:Addgene_70684). The *mCherry* coding sequence including three introns and flanking homology regions was ordered as gBlocks^®^ Gene Fragment from Integrated DNA Technologies™. All PCR products were purified using either the QIAquick^®^ PCR Purification Kit (Art. No. 28106, QIAGEN^®^) or the DNA Clean & Concentrator-5 Kit (Art. No. D4003, Zymo Research), eluted in sterile water and confirmed by agarose gel electrophoresis. For all epitope tag insertions, co-conversions and precise deletions, we ordered 4 nmole Ultramer^®^ DNA oligodeoxynucleotides from Integrated DNA Technologies™, which serves as linear, single-stranded DNA (ssODN) donor templates. All Ultramer^®^ DNA oligodeoxynucleotides were resuspended in sterile water. All linear DNA donor templates contained ∼35 bp homology regions (Paix et al., 2014, 2016), except the mCherry donor template, which contained 234 bp 5’ homology and 316 bp 3’ homology. All linear DNA donor templates are listed below.

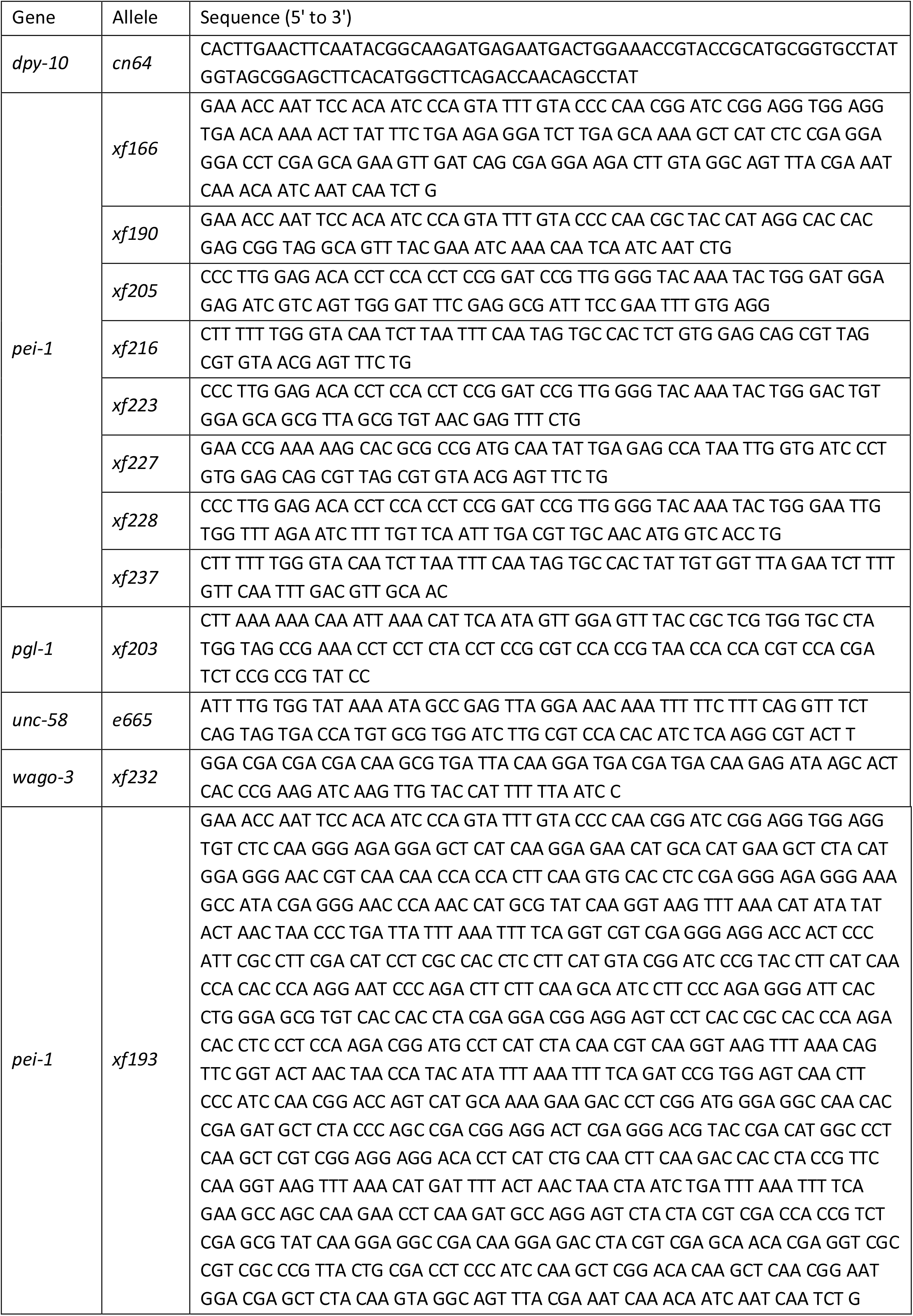
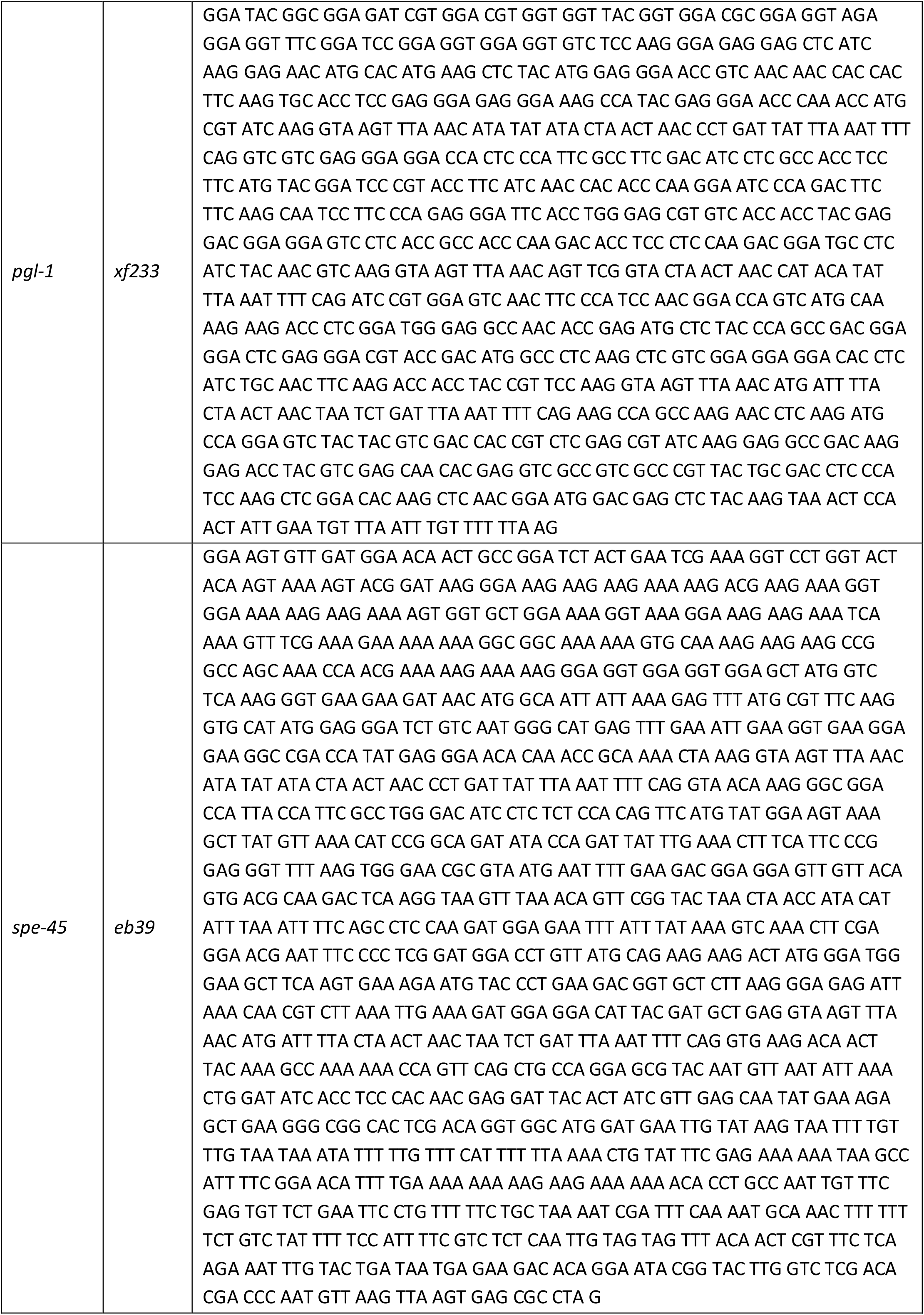

To generate the *mut-16* deletion allele, we injected animals with 50 ng/µl pJW1259, 30 ng/µl of each sgRNA(F+E), 10 ng/µl pGH8, 5 ng/µl pCFJ104, and 2.5 ng/µl pCFJ90. F1 animals expressing all three co-injection markers were selected for subsequent screening of *mut-16* deletion alleles. To insert *gfp::3xflag* sequences, injection mixes included 50 ng/µl pJW1259, 50 ng/µl sgRNA(F+E), 10 ng/µl SEC donor plasmid, 10 ng/µl pGH8, 5 ng/µl pCFJ104, and 2.5 ng/µl pCFJ90. Screening of F1 animals was performed as previously described (Dickinson et al., 2015). Every other CRISPR/Cas9-mediated genome editing was performed using either *dpy-10(cn64)* or *unc-58(e665)* co-conversion strategies (Arribere et al., 2014). To insert epitope tag or protospacer sequences, we injected 50 ng/µl Cas9 + sgRNA(F+E) (co-conversion), 50 ng/µl sgRNA(F+E) (gene of interest), 750 nM ssODN donor1 (co-conversion), and 750 nM ssODN donor2 (gene of interest). To insert a *mTagRfp-t* sequence in *pei-1* and *pgl-1*, we first transplanted the protospacer sequence used for the *dpy-10* co-conversion directly upstream of the respective stop codon to generate *d10*-entry strains (El Mouridi et al., 2017). These strains served as reference strains for the insertion of a *mTagRfp-t* sequence by injecting 50 ng/µl Cas9 + sgRNA(F+E) (*dpy-10* co-conversion), 1000 nM ssODN donor (*dpy-10* co-conversion), and 300 ng/µl linear, double-stranded DNA donor. Precise deletions in *pei-1* and *wago-3* were generated by injecting 50 ng/µl Cas9 + sgRNA(F+E) (co-conversion), 50 ng/µl of each sgRNA(F+E) (gene of interest), 750 nM ssODN donor1 (co-conversion), and 750 nM ssODN donor2 (gene of interest). To insert a *mCherry* sequence in *spe-45*, we injected a RNP mix containing 3.1 µM Alt-R^®^ S.p. Cas9 Nuclease 3NLS, 6.25 µM of each sgRNA targeting *spe-45*, 1.25 µM sgRNA targeting *dpy-10*, 500 nM ssODN (*dpy-10* co-conversion), and 120 ng/µl linear, double-stranded DNA donor (Paix et al., 2015). Alt-R^®^ S.p. Cas9 Nuclease 3NLS, Alt-R^®^ CRISPR-Cas9 tracrRNA and Alt-R^®^ CRISPR-Cas9 crRNA were purchased from Integrated DNA Technologies™. Unless otherwise stated, DNA injection mixes were injected in both gonad arms of five to 20 young adult N2 hermaphrodites maintained at 20°C. RNP injection mixes were injected in a single gonad arm of 40 young adult N2 hermaphrodites maintained at 20°C. Selected F1 progeny were screened for insertion or deletion by PCR. Successful editing events were confirmed by Sanger sequencing. All generated mutant strains were out-crossed at least two times prior to any further cross or analysis. All CRISPR/Cas9-generated alleles are listed below.

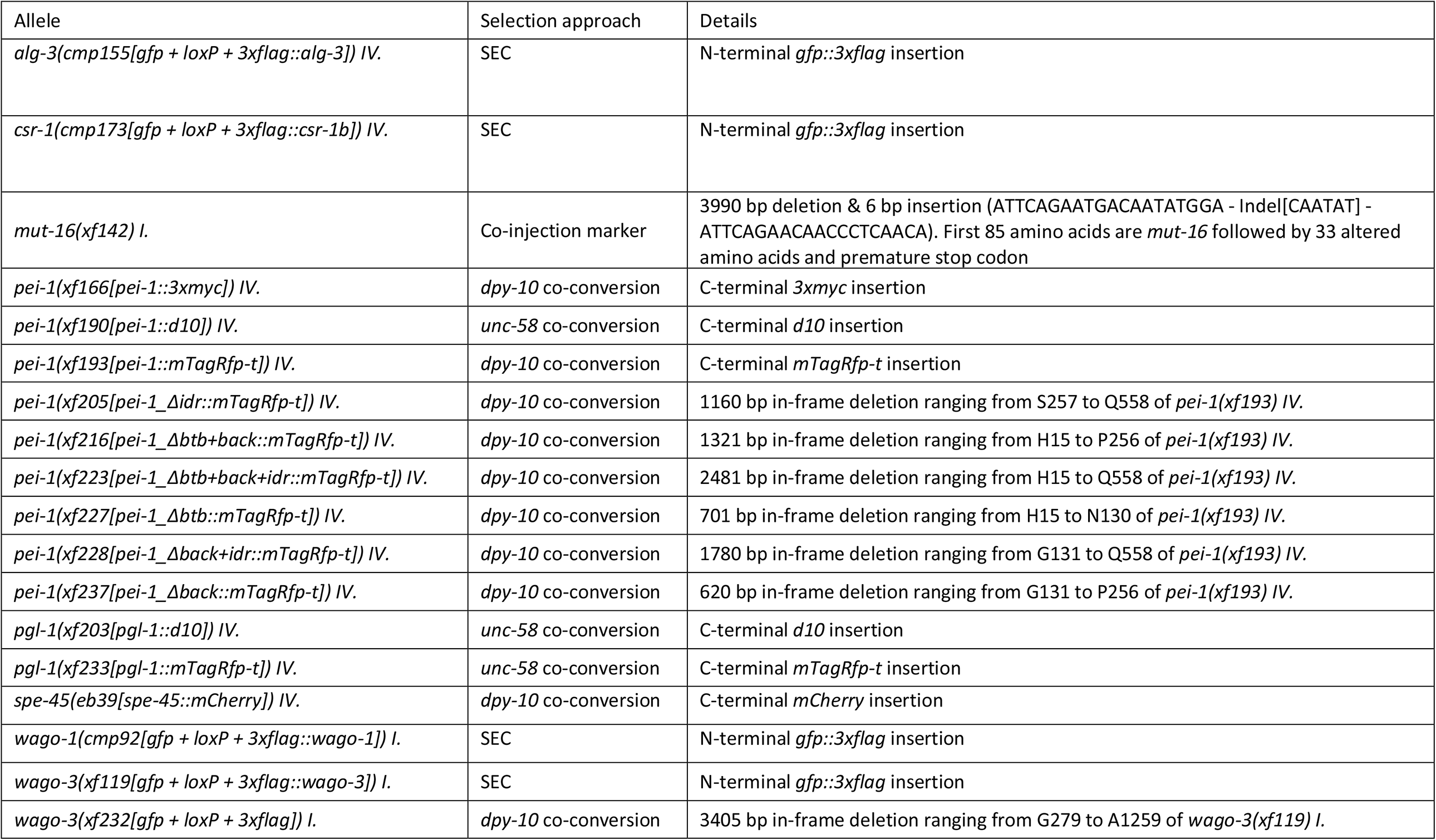

### Immunoprecipitation experiments

Unless otherwise stated, synchronized animals were cultured at 20°C until late-L4 stage, harvested with M9 buffer and frozen on dry ice in sterile water and 200 µl aliquots. Aliquots were thawed on ice, mixed with same volume of 2x lysis buffer (50 mM Tris HCl pH 7.5, 300 mM NaCl, 3 mM MgCl_2_, 2 mM DTT, 0.2 % Triton™ X-100, cOmplete™ Mini EDTA-free Protease Inhibitor Cocktail (Art. No. 11836170001, Roche)) and sonicated using a Bioruptor^®^ Plus device (Art. No. B01020001, Diagenode) (4°C, 10 cycles à 30 seconds ON and 30 seconds OFF). Following centrifugation for 10 min at 4°C and 21,000 xg, supernatants were carefully transferred into a fresh tube without taking any material from the pellet or lipid phase. Pellet fractions were washed three times in 1x lysis buffer and resuspended in 1x Novex™ NuPAGE™ LDS sample buffer (Art. No. NP0007, Invitrogen™) supplemented with 100 mM DTT. Total protein concentrations of soluble worm extracts were determined using the Pierce™ BCA™ Protein-Assay (Art. No. 23225, Thermo Scientific™) and an Infinite^®^ M200 Pro plate reader (Tecan). Extracts were diluted with 1x lysis buffer to reach 550 µl and a total protein concentration of 3 µg/µl. For each sample, 50 µl of this extract was added to 50 µl 1x Novex™ NuPAGE™ LDS sample buffer supplemented with 100 mM DTT and served as input control sample. For each immunoprecipitation (IP) experiment, 30 µl Novex™ DYNAL™ Dynabeads™ Protein G (Art. No. 10004D, Invitrogen™) were washed three times with 500 µl 1x wash buffer (25 mM Tris HCl pH 7.5, 150 mM NaCl, 1.5 mM MgCl_2_, 1 mM DTT, cOmplete™ Mini EDTA-free Protease Inhibitor Cocktail), combined with the remaining 500 µl extract and incubated with rotation for 1 h at 4°C. In the meantime, 8 µg antibody (Monoclonal ANTI-FLAG^®^ M2, Art. No. F3165, Sigma-Aldrich^®^ / Myc-Tag (9B11) Mouse mAb, Art. No. 2276, Cell Signaling Technology^®^) was conjugated to another 30 µl Novex™ DYNAL™ Dynabeads™ Protein G according to the manufacturer’s instructions. Extracts were separated from non-conjugated Dynabeads™, combined with antibody-conjugated Dynabeads™ and incubated with rotation for 2 h at 4°C. Following three washes with 500 µl 1x wash buffer, antibody-conjugated Dynabeads™ were resuspended in 25 µl 1.2x Novex™ NuPAGE™ LDS sample buffer supplemented with 120 mM DTT.

For RIP experiments, immunoprecipitations were performed as described above with the following modifications: i) adult animals were harvested, ii) soluble worm extract was diluted to 650 µl and a total protein concentration of 7 µg/µl, of which 150 µl served as input sample for later RNA extraction, iii) antibody-conjugated Dynabeads™ were resuspended in 50 µl nuclease-free water.

Immunoprecipitation experiments associated with mass spectrometry and small RNA sequencing were performed in quadruplicates and triplicates, respectively.

### Western Blot

Equal amounts of input samples (2 %) and IP samples (10 %) were adjusted to same volume with 1x Novex™ NuPAGE™ LDS sample buffer (Art. No. NP0007, Invitrogen™) supplemented with 100 mM DTT and incubated for 10 min at 95°C. Together with PageRuler™ Prestained Protein Ladder (10 to 180 kDa, Art. No. 26616, Thermo Scientific™), samples were separated on a Novex™ NuPAGE™ 4-12 % Bis-Tris Mini Protein Gel (Art. No. NP0323, Invitrogen™) in 1x Novex™ NuPAGE™ MOPS SDS Running Buffer (Art. No. NP0001, Invitrogen™) at 50 mA. Afterwards, proteins were transferred on an Immobilon™-P Membran (PVDF, 0.45 µm, Art. No. IPVH00010, Merck Millipore) for 16 h at 15 V using a Mini Trans-Blot^®^ Cell (Art. No. 1703930, Bio-Rad) and 1x NuPAGE™ Transfer Buffer (Art. No NP0006, Invitrogen™) supplemented with 20 % methanol. Following incubation in 1x PBS supplemented with 5 % skim milk and 0.05 % Tween^®^20 for 1 h, the PVDF membrane was cut according to the molecular weight of the proteins of interest. Each part was incubated in 1x PBS supplemented with 0.5 % skim milk, 0.05 % Tween^®^20 and the primary antibody (1:5,000 Monoclonal ANTI-FLAG^®^ M2, Art. No. F3165, Sigma-Aldrich^®^ / 1:1,000 Myc-Tag (9B11) Mouse mAb, Art. No. 2276, Cell Signaling Technology^®^ / 1:5,000 anti-Histone H3, Art. No. H0164, Sigma-Aldrich^®^) for 1 h, followed by three washes with 1x PBS supplemented with 0.05 % Tween^®^20 (hereinafter referred to as 0.05 % PBS-T) for 10 min each, one hour incubation in 0.05 % PBS-T supplemented with the secondary antibody (1:10,000 anti-mouse IgG, HRP-linked antibody, Art. No. 7076, Cell Signaling Technology^®^ / 1:10,000 anti-rabbit IgG, HRP-linked antibody, Art. No. 7074, Cell Signaling Technology^®^) and three final washes with 0.05 % PBS-T for 10 min each. Chemiluminescence detection was performed using Amersham™ ECL Select™ Western Blotting Detection Reagent (Art. No. RPN2235, GE Healthcare) and a ChemiDoc™ XRS+ System (Art. No. 1708265, Bio-Rad).

### Mass spectrometry and proteome comparison

IP samples resuspended in Novex™ NuPAGE™ LDS sample buffer (Art. No. NP0007, Invitrogen™) were incubated at 70°C for 10 min and separated on a Novex™ NuPAGE™ 4-12 % Bis-Tris Mini Protein Gel (Art. No. NP0321, Invitrogen™) in 1x Novex™ NuPAGE™ MOPS SDS Running Buffer (Art. No. NP0001, Invitrogen™) at 180 V for 10 min. After separation the samples were processed by in-gel digest as previously described (Kappei et al., 2013; Shevchenko et al., 2007). Following protein digest, the peptides were desalted using a C18 StageTip (Rappsilber et al., 2007). For measurement the digested peptides were separated on a 25 cm reverse-phase capillary (75 µm inner diameter) packed with Reprosil C18 material (Dr. Maisch GmbH). Elution was carried out along a two hour gradient of 2 to 40 % of a mixture of 80 % acetonitrile/0.5 % formic acid with the EASY-nLC 1000 system (Art. No. LC120, Thermo Scientific™). A Q Exactive™ Plus mass spectrometer (Thermo Scientific™) operated with a Top10 data-dependent MS/MS acquisition method per full scan was used for measurement (Bluhm et al., 2016). Processing of the obtained results was performed with the MaxQuant software, version 1.5.2.8 against the Wormbase protein database (version WS263) for quantitation (Cox and Mann, 2008). The processed data was visualized with R^®^ and R-Studio^®^ using in-house scripts. The mass spectrometry proteomics data have been deposited to the ProteomeXchange Consortium via the PRIDE (Perez-Riverol et al., 2019) partner repository with the dataset identifier PXD019099. Mitochondrial and spermatid proteins were described in previous proteomic studies (Jing et al., 2009; Ma et al., 2014). Genes encoding ribosomal proteins were obtained from Wormbase (version WS275). Comparison of protein lists was visualized with Intervene (Khan and Mathelier, 2017).

### RNA extraction, library preparation and sequencing

RNA of input and GFP::3xFLAG::WAGO-3 immunoprecipitation samples was extracted using TRIzol™ LS Reagenz (Art. No. 10296010, Invitrogen™) according to the manufacturer’s instructions, and resuspended in nuclease-free water. RNA quality and quantity was assessed using the Bioanalyzer RNA 6000 Nano Kit (Art. No. 5067-1511, Agilent Technologies) and Qubit™ RNA BR Assay Kit (Art. No. Q10210, Invitrogen™), respectively.

RNA 5’ Pyrophosphohydrolase (RppH) (Art. No. M0356S, New England BioLabs^®^) treatment was performed with a starting amount of 690 ng. After purification samples were quantified using the Qubit™ RNA HS Assay Kit (Art. No. Q32852, Invitrogen™). NGS library preparation was performed with NEXTFLEX^®^ Small RNA-Seq Kit v3 (Bioo Scientific^®^) following Step A to Step G of the manufacturer’s standard protocol (v16.06). Libraries were prepared with a starting amount ranging between 426 ng – 896 ng and amplified in 16 PCR cycles. Amplified libraries were purified by running an 8 % TBE gel and size-selected for 15 – 50 bp. Libraries were profiled in a High Sensitivity DNA Chip on a 2100 Bioanalyzer Instrument (Agilent Technologies) and quantified using the Qubit™ dsDNA HS Assay Kit (Art. No. Q32851, Invitrogen™), in a Qubit™ 2.0 Fluorometer (Invitrogen™). All samples were pooled in equimolar ratio and sequenced on one NextSeq 500/550 Flowcell, SR for 1x 84 cycles plus seven cycles for the index read. The accession number for the smRNA-seq data generated in this study is PRJNA629991 (https://dataview.ncbi.nlm.nih.gov/object/PRJNA629991?reviewer=72dsklt8i261719d2vq572tcq5).

### Read processing and mapping

Raw sequenced reads from high quality samples as assessed by FastQC were fed to Cutadapt (Martin, 2011) for adapter removal (-a TGGAATTCTCGGGTGCCAAGG -O 5 -m 26 -M 48) and low-quality reads were filtered out using the FASTX-Toolkit (fastq_quality_filter, -q 20 -p 100 -Q 33). Unique molecule identifiers (UMIs) were used to remove PCR duplicates via an in-house script and were subsequently removed using seqtk (trimfq-l 4 – b 4). Finally, reads shorter than 15 nt were removed with seqtk (seq -L 15). Reads were aligned to the *C. elegans* genome assembly WBcel235 using bowtie v1.2.2 (Langmead et al., 2009) (–phred33-quals –tryhard –best –strata –chunkmbs 256 -v 2 -M 1).

### Small RNA classification and target identification

A custom GTF-file was created by adding transposons retrieved from Wormbase (PRJNA13758.WS264) to the Ensembl reference WBcel235.84. Small RNAs classes were then defined as: 21U-RNAs, 21 nucleotide long mapped reads that map sense to annotated piRNA loci; 22G-RNAs, mapped reads of lengths 20-23 nucleotides, with no 5’ bias, and map antisense to protein-coding/pseudogenes/lincRNA/transposons; 26G-RNAs, mapped reads 26 nucleotides long and map antisense to annotated protein-coding/pseudogenes/lincRNA; miRNAs are 20-24 nt reads mapping sense to annotated miRNA loci; finally all mapped reads longer than 26 nucleotides were classed in a separate group. Read filtering was done with a python script (https://github.com/adomingues/filterReads/blob/master/filterReads/filterSmallRNAclasses.py) based on pysam v0.8.1 / htslib (Li et al., 2009), in combination with BEDTools intersect (Quinlan and Hall, 2010).

22G reads mapping to features in the custom annotation were counted using htseq-count v0.9.0 (Anders et al., 2015) (-s no -m intersection-nonempty). Differential targeting analysis was carried out in R^®^ using DESeq2 (Love et al., 2014) with a strict cut-off for the adjusted p-value of 0.01. A cut-off for fold-change (IP versus input) was determined by fitting a Gaussian to the fold-change-distribution of reads mapping to miRNA, which are known not to bind WAGOs, and choosing the value that corresponds to a false discovery rate (FDR) of 5% for this RNA species; here log2-fold-change > 1.3.

Protein-coding target genes of WAGO-3 were compared to: i) protein-coding target genes of CSR-1 (Claycomb et al., 2009), ii) protein-coding target genes of siRNAs downregulated in *mut-16* mutant animals (Phillips et al., 2014), and iii) protein-coding target genes of sperm-derived 22G RNAs (Stoeckius et al., 2014). To determine germline expression, protein-coding target genes of WAGO-3 were compared to lists of genes expressed in the *C. elegans* germline of either *fem-3* or *fog-2* mutant animals (Ortiz et al., 2014).

### 22G RNA coverage on protein-coding genes

Coverage of 22Gs along targeted protein coding genes was visualized by i) creating bigwig tracks normalized to mapped non-structural reads (rRNA/tRNA/snoRNA/snRNA) * 1 million (RPM) using a combination of BEDTools (genomeCoverageBed -bg -scale -split) (Quinlan and Hall, 2010) followed by bedGraphToBigWig; ii) log_2_(IP/input) normalized tracks were created with deepTools v2.4.2 (Ramírez et al., 2014) (bigwigCompare –binSize 10 –ratio log2); iii) coverage for each gene was determine with deepTools (computeMatrix scale-regions --metagene --missingDataAsZero -b 250 -a 250 -- regionBodyLength 2000 --binSize 50 --averageTypeBins median); and plots generated with plotProfile (--plotType se --averageType mean --perGroup) to scale and visualize 22G abundance along targeted genes.

Reads mapping to intronic, exonic, or untranslated regions were counted using a custom Python script. Reads mapping at exon-intron junctions were counted as 0.5 intronic and 0.5 exonic regardless of the spanned region.

### Microscopy

For L4 larvae, adults and males, 20 – 30 animals were washed in a drop of 100 µl M9 buffer and subsequently transferred to a drop of 50 µl M9 buffer supplemented with 40 mM sodium azide on a coverslip. After 15 to 30 min, excess buffer was removed and a glass slide containing a freshly made agarose pad (2 % (w/v) in water) was placed on top of the coverslip. For imaging embryos, adult hermaphrodites were washed and dissected in M9 buffer before mounting. To image sperm, L4 males were singled from hermaphrodites, grown over night, washed and dissected in SMG buffer (50 mM HEPES pH 7.5, 50 mM NaCl, 25 mM KCl, 5 mM CaCl_2_, and 1 mM MgSO_4_, 10 mM glucose) by cutting near the vas deferens. Animals and sperm were immediately imaged using a TCS SP5 Leica confocal microscope equipped with a HCX PL APO 63x water objective (NA 1.2) or HCX PL APO CS 40x oil objective (NA 1.3). Fluorescence emission was detected by either photomultiplier tubes (PMTs) or hybrid detectors (HyDs). Depending on the experiment, SMG buffer was supplemented with 1:2,000 Hoechst33342 (Art. No. H1399, Invitrogen™), 200 nM MitoTracker^®^ Green FM (Art. No. M7514, Invitrogen™) or 1,6-hexanediol (Art. No. 240117, Sigma-Aldrich^®^), and sperm were imaged after 30 min incubation. To score the expression of a germline-specific mCherry::H2B transgene, we used a Leica DM6000 B research microscope equipped with a HC PL Fluotar 20x dry objective (NA 0.5). Images were processed using Fiji and the following figures were deconvolved using the Huygens Remote Manager v3.6: Figures 2D-2H; 6A; S2; S3; S4D-4F; S6A.

Time series of spermatocytes expressing GFP::3xFLAG::WAGO-3 were acquired with a fluorescence spinning disk confocal microscope (SDCM) from Visitron Systems (VisiSope 5Elements) based on a Nikon Ti-2E stand and a spinning disk from Yokogawa (CSU-W, 50 µm pinhole) controlled by the VisiView^®^ software. The microscope was equipped with a 60x plan apochromatic water immersion objective (CFI Plan Apo VC, NA 1.2), a twofold magnification lens in front of the sCMOS camera (BSI, Photometrics), and a stage-top incubation chamber for live imaging (20°C, ambient CO_2_). The sample was excited by an argon laser at λ_ex_ = 488 nm (200 mW, power set to 20 %) and the emission was detected in a range of λ_em_ = 500 - 550 nm (ET525/50m, Chroma).

### FRAP

FRAP measurements were performed on a TCS SP5 Leica confocal microscope equipped with a FRAP-booster and a HCX PL APO 63x water objective (NA 1.2). An entire granule was bleached in a fixed region of interest (ROI) (0.9 µm diameter), while two additional control ROIs of same size were used to detect fluorescence emission of an unbleached granule and background signal, respectively. Five pre-bleach frames were recorded (5x 0.374 s/frame), followed by two bleach frames (2x 0.374 s/frame), and 3 sets of post-bleach frames (10x 0.5 s/frame, 10x 5 s/frame, 15x 10 s/frame). Data analysis including full scale normalization and curve fitting using a double term exponential equation was performed using EasyFRAP-web (Koulouras et al., 2018)

## REFERENCES

de Albuquerque, B.F.M., Placentino, M., and Ketting, R.F. (2015). Maternal piRNAs Are Essential for Germline Development following De Novo Establishment of Endo-siRNAs in Caenorhabditis elegans. Dev. Cell 34, 448–456.

Alcazar, R.M., Lin, R., and Fire, A.Z. (2008). Transmission dynamics of heritable silencing induced by double-stranded RNA in Caenorhabditis elegans. Genetics 180, 1275–1288.

Anders, S., Pyl, P.T., and Huber, W. (2015). HTSeq-A Python framework to work with high-throughput sequencing data. Bioinformatics 31, 166–169.

Arribere, J.A., Bell, R.T., Fu, B.X.H., Artiles, K.L., Hartman, P.S., and Fire, A.Z. (2014). Efficient marker-free recovery of custom genetic modifications with CRISPR/Cas9 in caenorhabditis elegans. Genetics 198, 837–846.

Bagijn, M.P., Goldstein, L.D., Sapetschnig, A., Weick, E., Bouasker, S., Lehrbach, N.J., Simard, M.J., and Miska, E.A. (2012). Function, Targets, and Evolution of Caenorhabditis elegans piRNAs. Science (80-.). 337, 574–578.

Banani, S.F., Lee, H.O., Hyman, A.A., and Rosen, M.K. (2017). Biomolecular condensates: Organizers of cellular biochemistry. Nat. Rev. Mol. Cell Biol. 18, 285–298.

Batista, P.J., Ruby, J.G., Claycomb, J.M., Chiang, R., Fahlgren, N., Kasschau, K.D., Chaves, D.A., Gu, W., Vasale, J.J., Duan, S., et al. (2008). PRG-1 and 21U-RNAs Interact to Form the piRNA Complex Required for Fertility in C. elegans. Mol. Cell 31, 67–78.

Bluhm, A., Casas-Vila, N., Scheibe, M., and Butter, F. (2016). Reader interactome of epigenetic histone marks in birds. Proteomics 16, 427–436.

Bošković, A., and Rando, O.J. (2018). Transgenerational epigenetic inheritance. Annu. Rev. Genet. 52, 21–41.

Brangwynne, C.P., Eckmann, C.R., Courson, D.S., Rybarska, A., Hoege, C., Gharakhani, J., Jülicher, F., and Hyman, A.A. (2009). Germline P granules are liquid droplets that localize by controlled dissolution/condensation. Science (80-.). 324, 1729–1732.

Brenner, S. (1974). The genetics of Caenorhabditis elegans. Genetics 77, 71–94.

Buckley, B.A., Burkhart, K.B., Gu, S.G., Spracklin, G., Kershner, A., Fritz, H., Kimble, J., Fire, A., and Kennedy, S. (2012). A nuclear Argonaute promotes multigenerational epigenetic inheritance and germline immortality. Nature 489, 447–451.

Castel, S.E., and Martienssen, R.A. (2013). RNA interference in the nucleus: Roles for small RNAs in transcription, epigenetics and beyond. Nat. Rev. Genet. 14, 100–112.

Chen, B., Gilbert, L.A., Cimini, B.A., Schnitzbauer, J., Zhang, W., Li, G.W., Park, J., Blackburn, E.H., Weissman, J.S., Qi, L.S., et al. (2013). Dynamic imaging of genomic loci in living human cells by an optimized CRISPR/Cas system. Cell 155, 1479–1491.

Chiu, J., March, P.E., Lee, R., and Tillett, D. (2004). Site-directed, Ligase-Independent Mutagenesis (SLIM): a single-tube methodology approaching 100% efficiency in 4 h. Nucleic Acids Res. 32.

Chiu, J., Tillett, D., Dawes, I.W., and March, P.E. (2008). Site-directed, Ligase-Independent Mutagenesis (SLIM) for highly efficient mutagenesis of plasmids greater than 8kb. J. Microbiol. Methods 73, 195–198.

Claycomb, J.M., Batista, P.J., Pang, K.M., Gu, W., Vasale, J.J., van Wolfswinkel, J.C., Chaves, D.A., Shirayama, M., Mitani, S., Ketting, R.F., et al. (2009). The Argonaute CSR-1 and Its 22G-RNA Cofactors Are Required for Holocentric Chromosome Segregation. Cell 139, 123–134.

Collins, T., Stone, J.R., and Williams, A.J. (2001). All in the Family: the BTB/POZ, KRAB, and SCAN Domains. Mol. Cell. Biol. 21, 3609–3615.

Conine, C.C., Batista, P.J., Gu, W., Claycomb, J.M., Chaves, D.A., Shirayama, M., and Mello, C.C. (2010). Argonautes ALG-3 and ALG-4 are required for spermatogenesis-specific 26G-RNAs and thermotolerant sperm in Caenorhabditis elegans. Proc. Natl. Acad. Sci. U. S. A. 107, 3588–3593.

Conine, C.C., Moresco, J.J., Gu, W., Shirayama, M., Conte, D., Yates, J.R., and Mello, C.C. (2013). Argonautes promote male fertility and provide a paternal memory of germline gene expression in C. Elegans. Cell 155, 1532–1544.

Cox, J., and Mann, M. (2008). MaxQuant enables high peptide identification rates, individualized p.p.b.-range mass accuracies and proteome-wide protein quantification. Nat. Biotechnol. 26, 1367–1372.

Das, P.P., Bagijn, M.P., Goldstein, L.D., Woolford, J.R., Lehrbach, N.J., Sapetschnig, A., Buhecha, H.R., Gilchrist, M.J., Howe, K.L., Stark, R., et al. (2008). Piwi and piRNAs Act Upstream of an Endogenous siRNA Pathway to Suppress Tc3 Transposon Mobility in the Caenorhabditis elegans Germline. Mol. Cell 31, 79–90.

Dickinson, D.J., Ward, J.D., Reiner, D.J., and Goldstein, B. (2013). Engineering the Caenorhabditis elegans genome using Cas9-triggered homologous recombination. Nat. Methods 10, 1028–1034.

Dickinson, D.J., Pani, A.M., Heppert, J.K., Higgins, C.D., and Goldstein, B. (2015). Streamlined genome engineering with a self-excising drug selection cassette. Genetics 200, 1035–1049.

Drozdetskiy, A., Cole, C., Procter, J., and Barton, G.J. (2015). JPred4: a protein secondary structure prediction server. Nucleic Acids Res. 43, W389–W394.

Ellis, R.E., and Stanfield, G.M. (2014). The regulation of spermatogenesis and sperm function in nematodes. Semin. Cell Dev. Biol. 29, 17–30.

Fabig, G., Schwarz, A., Striese, C., Laue, M., and Müller-Reichert, T. (2019). In situ analysis of male meiosis in C. elegans. In Methods in Cell Biology, pp. 119–134.

Frøkjær-Jensen, C., Wayne Davis, M., Hopkins, C.E., Newman, B.J., Thummel, J.M., Olesen, S.P., Grunnet, M., and Jorgensen, E.M. (2008). Single-copy insertion of transgenes in Caenorhabditis elegans. Nat. Genet. 40, 1375–1383.

Ge, D.T., Wang, W., Tipping, C., Gainetdinov, I., Weng, Z., and Zamore, P.D. (2019). The RNA-Binding ATPase, Armitage, Couples piRNA Amplification in Nuage to Phased piRNA Production on Mitochondria. Mol. Cell 74, 982-995.e6.

Gjerstorff, M.F., Rösner, H.I., Pedersen, C.B., Greve, K.B. V., Schmidt, S., Wilson, K.L., Mollenhauer, J., Besir, H., Poulsen, F.M., Møllegaard, N.E., et al. (2012). GAGE Cancer-Germline Antigens Are Recruited to the Nuclear Envelope by Germ Cell-Less (GCL). PLoS One 7, e45819.

Gleason, E.J., Lindsey, W.C., Kroft, T.L., Singson, A.W., and L’Hernault, S.W. (2006). Spe-10 encodes a DHHC-CRD zinc-finger membrane protein required for endoplasmic reticulum/golgi membrane morphogenesis during Caenorhabditis elegans spermatogenesis. Genetics 172, 145–158.

Gold, V.A., Chroscicki, P., Bragoszewski, P., and Chacinska, A. (2017). Visualization of cytosolic ribosomes on the surface of mitochondria by electron cryo-tomography. EMBO Rep. 18, 1786–1800.

Grishok, A., Tabara, H., and Mello, C.C. (2000). Genetic requirements for inheritance of RNAi in C. elegans. Science (80-.). 287, 2494–2497.

Gu, W., Shirayama, M., Conte, D., Vasale, J., Batista, P.J., Claycomb, J.M., Moresco, J.J., Youngman, E.M., Keys, J., Stoltz, M.J., et al. (2009). Distinct Argonaute-Mediated 22G-RNA Pathways Direct Genome Surveillance in the C. elegans Germline. Mol. Cell 36, 231–244.

Haeussler, M., Schönig, K., Eckert, H., Eschstruth, A., Mianné, J., Renaud, J.-B., Schneider-Maunoury, S., Shkumatava, A., Teboul, L., Kent, J., et al. (2016). Evaluation of off-target and on-target scoring algorithms and integration into the guide RNA selection tool CRISPOR. Genome Biol. 17, 148.

Hanazawa, M., Yonetani, M., and Sugimoto, A. (2011). PGL proteins self associate and bind RNPs to mediate germ granule assembly in C. elegans. J. Cell Biol. 192, 929–937.

Hofweber, M., and Dormann, D. (2019). Friend or foe-Post-translational modifications as regulators of phase separation and RNP granule dynamics. J. Biol. Chem. 294, 7137–7150.

Huerta-Cepas, J., Szklarczyk, D., Forslund, K., Cook, H., Heller, D., Walter, M.C., Rattei, T., Mende, D.R., Sunagawa, S., Kuhn, M., et al. (2016). eggNOG 4.5: a hierarchical orthology framework with improved functional annotations for eukaryotic, prokaryotic and viral sequences. Nucleic Acids Res. 44, D286–D293.

Hutvagner, G., and Simard, M.J. (2008). Argonaute proteins: Key players in RNA silencing. Nat. Rev. Mol. Cell Biol. 9, 22–32.

Hyman, A.A., Weber, C.A., and Jülicher, F. (2014). Liquid-Liquid Phase Separation in Biology. Annu. Rev. Cell Dev. Biol. 30, 39–58.

Jing, L., Tanxi, C., Peng, W., Ziyou, C., Xiulan, C., Junjie, H., Zhensheng, X., Peng, X., Linan, S., Pingsheng, L., et al. (2009). Proteomic analysis of mitochondria from Caenorhabditis elegans. Proteomics 9, 4539–4553.

Kappei, D., Butter, F., Benda, C., Scheibe, M., Draškovič, I., Stevense, M., Novo, C.L., Basquin, C., Araki, M., Araki, K., et al. (2013). HOT1 is a mammalian direct telomere repeat-binding protein contributing to telomerase recruitment. EMBO J. 32, 1681–1701.

Kawasaki, I., Shim, Y.H., Kirchner, J., Kaminker, J., Wood, W.B., and Strome, S. (1998). PGL-1, a predicted RNA-binding component of germ granules, is essential for fertility in C. elegans. Cell 94, 635–645.

Kawasaki, I., Amiri, A., Fan, Y., Meyer, N., Dunkelbarger, S., Motohashi, T., Karashima, T., Bossinger, O., and Strome, S. (2004). The PGL family proteins associate with germ granules and function redundantly in Caenorhabditis elegans germline development. Genetics 167, 645–661.

Kelleher, J.F., Mandell, M.A., Moulder, G., Hill, K.L., L’Hernault, S.W., Barstead, R., and Titus, M.A. (2000). Myosin VI is required for asymmetric segregation of cellular components during C. elegans spermatogenesis. Curr. Biol. 10, 1489–1496.

Khan, A., and Mathelier, A. (2017). Intervene: A tool for intersection and visualization of multiple gene or genomic region sets. BMC Bioinformatics 18, 1–8.

Kleiman, S.E., Yogev, L., Gal-Yam, E.N., Hauser, R., Gamzu, R., Botchan, A., Paz, G., Yavetz, H., Maymon, B.B.S., Schreiber, L., et al. (2003). Reduced Human Germ Cell-Less (HGCL) Expression in Azoospermic Men with Severe Germinal Cell Impairment. J. Androl. 24, 670–675.

Kotaja, N., and Sassone-corsi, P. (2007). The chromatoid body: a germ-cell-specific RNA-processing centre. Nat. Rev. Mol. Cell Biol. 8, 85–90.

Koulouras, G., Panagopoulos, A., Rapsomaniki, M.A., Giakoumakis, N.N., Taraviras, S., and Lygerou, Z. (2018). EasyFRAP-web: A web-based tool for the analysis of fluorescence recovery after photobleaching data. Nucleic Acids Res. 46, W467–W472.

Kroschwald, S., Maharana, S., and Simon, A. (2017). Hexanediol: a chemical probe to investigate the material properties of membrane-less compartments. Matters 1–7.

Langmead, B., Trapnell, C., Pop, M., and Salzberg, S.L. (2009). Ultrafast and memory-efficient alignment of short DNA sequences to the human genome. Genome Biol. 10.

Lee, H.C., Gu, W., Shirayama, M., Youngman, E., Conte, D., and Mello, C.C. (2012). C. elegans piRNAs mediate the genome-wide surveillance of germline transcripts. Cell 150, 78–87.

Lev, I., Toker, I.A., Mor, Y., Nitzan, A., Weintraub, G., Antonova, O., Bhonkar, O., Ben Shushan, I., Seroussi, U., Claycomb, J.M., et al. (2019). Germ Granules Govern Small RNA Inheritance. Curr. Biol. 29, 2880-2891.e4.

Li, H., Handsaker, B., Wysoker, A., Fennell, T., Ruan, J., Homer, N., Marth, G., Abecasis, G., and Durbin, R. (2009). The Sequence Alignment/Map format and SAMtools. Bioinformatics 25, 2078–2079.

Love, M.I., Huber, W., and Anders, S. (2014). Moderated estimation of fold change and dispersion for RNA-seq data with DESeq2. Genome Biol. 15, 1–21.

Luteijn, M.J., and Ketting, R.F. (2013). PIWI-interacting RNAs: From generation to transgenerational epigenetics. Nat. Rev. Genet. 14, 523–534.

Ma, X., Zhu, Y., Li, C., Xue, P., Zhao, Y., Chen, S., Yang, F., and Miao, L. (2014). Characterisation of Caenorhabditis elegans sperm transcriptome and proteome. BMC Genomics 15, 1–13.

Mao, H., Zhu, C., Zong, D., Weng, C., Yang, X., Huang, H., Liu, D., Feng, X., and Guang, S. (2015). The Nrde Pathway Mediates Small-RNA-Directed Histone H3 Lysine 27 Trimethylation in Caenorhabditis elegans. Curr. Biol. 25, 2398–2403.

Martin, M. (2011). Cutadapt removes adapter sequences from high-throughput sequencing reads. EMBnet.Journal 17, 10.

El Mouridi, S., Lecroisey, C., Tardy, P., Mercier, M., Leclercq-Blondel, A., Zariohi, N., and Boulin, T. (2017). Reliable CRISPR/Cas9 genome engineering in Caenorhabditis elegans using a single efficient sgRNA and an easily recognizable phenotype. G3 Genes, Genomes, Genet. 7, 1429–1437.

Munafò, M., Manelli, V., Falconio, F.A., Sawle, A., Kneuss, E., Eastwood, E.L., Seah, J.W.E., Czech, B., and Hannon, G.J. (2019). Daedalus and gasz recruit armitage to mitochondria, bringing piRNA precursors to the biogenesis machinery. Genes Dev. 33, 844–856.

Ortiz, M.A., Noble, D., Sorokin, E.P., and Kimble, J. (2014). A New Dataset of Spermatogenic vs . Oogenic Transcriptomes in the Nematode Caenorhabditis elegans. G3 Genes, Genomes, Genet. 4, 1765–1772.

Ozata, D.M., Gainetdinov, I., Zoch, A., O’Carroll, D., and Zamore, P.D. (2019). PIWI-interacting RNAs: small RNAs with big functions. Nat. Rev. Genet. 20, 89–108.

Paix, A., Wang, Y., Smith, H.E., Lee, C.Y.S., Calidas, D., Lu, T., Smith, J., Schmidt, H., Krause, M.W., and Seydoux, G. (2014). Scalable and versatile genome editing using linear DNAs with microhomology to Cas9 sites in Caenorhabditis elegans. Genetics 198, 1347–1356.

Paix, A., Folkmann, A., Rasoloson, D., and Seydoux, G. (2015). High efficiency, homology-directed genome editing in C.elegans using CRISPR/Cas9 ribonucleoprotein complex. Genetics.

Paix, A., Schmidt, H., and Seydoux, G. (2016). Cas9-assisted recombineering in C. elegans: Genome editing using in vivo assembly of linear DNAs. Nucleic Acids Res. 44, e128.

Perez-Riverol, Y., Csordas, A., Bai, J., Bernal-Llinares, M., Hewapathirana, S., Kundu, D.J., Inuganti, A., Griss, J., Mayer, G., Eisenacher, M., et al. (2019). The PRIDE database and related tools and resources in 2019: Improving support for quantification data. Nucleic Acids Res. 47, D442–D450.

Perez, M.F., and Lehner, B. (2019). Intergenerational and transgenerational epigenetic inheritance in animals. Nat. Cell Biol. 21, 143–151.

Peters, L., and Meister, G. (2007). Argonaute Proteins: Mediators of RNA Silencing. Mol. Cell 26, 611–623.

Phillips, C.M., Montgomery, T.A., Breen, P.C., and Ruvkun, G. (2012). MUT-16 promotes formation of perinuclear Mutator foci required for RNA silencing in the C. elegans germline. Genes Dev. 26, 1433–1444.

Phillips, C.M., Montgomery, B.E., Breen, P.C., Roovers, E.F., Rim, Y.S., Ohsumi, T.K., Newman, M.A., Van Wolfswinkel, J.C., Ketting, R.F., Ruvkun, G., et al. (2014). MUT-14 and SMUT-1 DEAD box RNA helicases have overlapping roles in germline RNAi and endogenous siRNA formation. Curr. Biol. 24, 839–844.

Phillips, C.M., Brown, K.C., Montgomery, B.E., Ruvkun, G., and Montgomery, T.A. (2015). PiRNAs and piRNA-Dependent siRNAs Protect Conserved and Essential C. elegans Genes from Misrouting into the RNAi Pathway. Dev. Cell 34, 457–465.

Putnam, A., Cassani, M., Smith, J., and Seydoux, G. (2019). A gel phase promotes condensation of liquid P granules in Caenorhabditis elegans embryos. Nat. Struct. Mol. Biol. 26, 220–226.

Quinlan, A.R., and Hall, I.M. (2010). BEDTools: A flexible suite of utilities for comparing genomic features. Bioinformatics 26, 841–842.

Ramírez, F., Dündar, F., Diehl, S., Grüning, B.A., and Manke, T. (2014). DeepTools: A flexible platform for exploring deep-sequencing data. Nucleic Acids Res. 42, 187–191.

Rappsilber, J., Mann, M., and Ishihama, Y. (2007). Protocol for micro-purification, enrichment, pre-fractionation and storage of peptides for proteomics using StageTips. Nat. Protoc. 2, 1896–1906.

Ren, J., Wen, L., Gao, X., Jin, C., Xue, Y., and Yao, X. (2008). CSS-Palm 2.0: An updated software for palmitoylation sites prediction. Protein Eng. Des. Sel. 21, 639–644.

Robert, V.P. V, Sijen, T., van Wolfswinkel, J., and Plasterk, R.H.A. (2005). Chromatin and RNAi factors protect the C. elegans germline against repetitive sequences. Genes Dev. 19, 782–787.

Sato, K., and Sato, M. (2017). Multiple ways to prevent transmission of paternal mitochondrial DNA for maternal inheritance in animals. J. Biochem. 162, 247–253.

Schweinsberg, P.J., and Grant, B.D. (2013). C. elegans gene transformation by microparticle bombardment. In WormBook, pp. 1–10.

Shevchenko, A., Tomas, H., Havliš, J., Olsen, J. V., and Mann, M. (2007). In-gel digestion for mass spectrometric characterization of proteins and proteomes. Nat. Protoc. 1, 2856–2860.

Spike, C.A., Bader, J., Reinke, V., and Strome, S. (2008). DEPS-1 promotes P-granule assembly and RNA interference in C. elegans germ cells. Development 135, 983–993.

Stoeckius, M., Grün, D., and Rajewsky, N. (2014). Paternal RNA contributions in the Caenorhabditis elegans zygote. EMBO J. 33, 1740–1750.

Stogios, P.J., and Privé, G.G. (2004). The BACK domain in BTB-kelch proteins. Trends Biochem. Sci. 29, 634–637.

Tabaczar, S., Czogalla, A., Podkalicka, J., Biernatowska, A., and Sikorski, A.F. (2017). Protein palmitoylation: Palmitoyltransferases and their specificity. Exp. Biol. Med. 242, 1150–1157.

Updike, D., and Strome, S. (2010). P granule assembly and function in Caenorhabditis elegans germ cells. J. Androl. 31, 53–60.

Vacic, V., Uversky, V.N., Dunker, A.K., and Lonardi, S. (2007). Composition Profiler: a tool for discovery and visualization of amino acid composition differences. BMC Bioinformatics 8, 211.

Vastenhouw, N.L., Fischer, S.E.J., Robert, V.J.P., Thijssen, K.L., Fraser, A.G., Kamath, R.S., Ahringer, J., and Plasterk, R.H.A. (2003). A genome-wide screen identifies 27 genes involved in transposon silencing in C. elegans. Curr. Biol. 13, 1311–1316.

Voronina, E., Seydoux, G., Sassone-Corsi, P., and Nagamori, I. (2011). RNA granules in germ cells. Cold Spring Harb. Perspect. Biol. 3.

Wan, G., Fields, B.D., Spracklin, G., Shukla, A., Phillips, C.M., and Kennedy, S. (2018). Spatiotemporal regulation of liquid-like condensates in epigenetic inheritance. Nature 557, 679–683.

Wang Lv, Guo, and Yuan (2020). Mitochondria Associated Germinal Structures in Spermatogenesis: piRNA Pathway Regulation and Beyond. Cells 9, 399.

Wang, J., Choi, J.-M., Holehouse, A.S., Lee, H.O., Zhang, X., Jahnel, M., Maharana, S., Lemaitre, R., Pozniakovsky, A., Drechsel, D., et al. (2018). A Molecular Grammar Governing the Driving Forces for Phase Separation of Prion-like RNA Binding Proteins. Cell 174, 688-699.e16.

Ward, J.D. (2014). Rapid and precise engineering of the caenorhabditis elegans genome with lethal mutation co-conversion and inactivation of NHEJ repair. Genetics 199, 363–377.

Ward, S., Argon, Y., and Nelson, G.A. (1981). Sperm morphogenesis in wild-type and fertilization-defective mutants of Caenorhabditis elegans. J. Cell Biol. 91, 26–44.

Xie, Y., Zheng, Y., Li, H., Luo, X., He, Z., Cao, S., Shi, Y., Zhao, Q., Xue, Y., Zuo, Z., et al. (2016). GPS-Lipid: a robust tool for the prediction of multiple lipid modification sites. Sci. Rep. 6, 28249.

Xu, F., Guang, S., and Feng, X. (2018). Distinct nuclear and cytoplasmic machineries cooperatively promote the inheritance of RNAi in Caenorhabditis elegans. Biol. Cell 110, 217–224.

Yigit, E., Batista, P.J., Bei, Y., Pang, K.M., Chen, C.-C.G., Tolia, N.H., Joshua-Tor, L., Mitani, S., Simard, M.J., and Mello, C.C. (2006). Analysis of the C. elegans Argonaute Family Reveals that Distinct Argonautes Act Sequentially during RNAi. Cell 127, 747–757.

Yushin, V. V., and Malakhov, V. V. (2014). The origin of nematode sperm: Progenesis at the cellular level. Russ. J. Mar. Biol. 40, 71–81.

Zhang, C., Montgomery, T.A., Gabel, H.W., Fischer, S.E.J., Phillips, C.M., Fahlgren, N., Sullivan, C.M., Carrington, J.C., and Ruvkun, G. (2011). mut-16 and other mutator class genes modulate 22G and 26G siRNA pathways in Caenorhabditis elegans. Proc. Natl. Acad. Sci. U. S. A. 108, 1201–1208.

Zhou, L., Canagarajah, B., Zhao, Y., Baibakov, B., Tokuhiro, K., Maric, D., and Dean, J. (2017). BTBD18 Regulates a Subset of piRNA-Generating Loci through Transcription Elongation in Mice. Dev. Cell 40, 453-466.e5.

