## Supplemental text and Figures for "A membrane-associated condensate drives paternal epigenetic inheritance in *C. elegans*"

**Supplemental figure legends**

**Figure S1. Related to Figure 1.**

A. Read length distribution and first nucleotide bias of indicated small RNA libraries.

B. Classification of WAGO-3-targeted protein-coding genes according to germline expression (left panel) or gamete-specific expression (right panel). Statistical significance was tested with a Chi-square test (***: $p \leq 0.001$, ns: $p > 0.05$).

C. Dissection of 22G RNA reads mapping to protein-coding genes.

D. Metagene analysis of 22G RNA reads mapping to protein-coding target genes. TSS – transcription start site, TES – transcription end site. Shading around each line represents the standard error of each bin.

**Figure S2. Related to Figure 2.**

A-I: Confocal micrographs of an adult hermaphrodite (A), embryo (B), male (C) and various late-L4 stage hermaphrodites (D-I) expressing indicated proteins. Zooms show perinuclear co-localization of GFP::3xFLAG::WAGO-3 and PGL-1::mTagRFP-T in meiotic (I) and primordial (II) germ cells. Panel A and B show confocal maximum intensity projections. Scale bars: 20 µm (A, adult), 20 µm (B, embryo), 20 µm (C), 10 µm (A, zoom), 4 µm (B, zoom), 10 µm (D-I).

**Figure S3. Related to Figure 2 and 5.**

Confocal micrographs showing spermatogenesis of late-L4 stage hermaphrodites expressing PEI-1::mTagRFP-T in indicated mutants. sc – spermatocyte, rb – residual body, st – spermatid. Scale bars: 10 µm.

**Figure S4. Related to Figure 4.**

A. Schematic representation of the transcriptional GFP reporter from the endogenous *wago-3* locus.

B. Confocal micrograph of a L4 stage hermaphrodite expressing GFP::3xFLAG from the endogenous *wago-3* locus. Insert shows presence of GFP::3xFLAG within the spermatheca. sc – spermatocyte, rb – residual body, st – spermatid. Scale bars: 20 µm (L4 gonad), 10 µm (insert, spermatheca).

C. Confocal maximum intensity projections of isolated male-derived spermatocytes and budding spermatids expressing indicated proteins. White arrow heads mark budding spermatids, red arrows indicate residual bodies. Hoechst33342 was used to stain DNA. Scale bars: 4 µm.

D. Confocal Z-stack of a spermatocyte expressing PEI-1_ΔBTB+BACK+IDR::mTagRFP-T as shown in Figure 4C. Z-size: 125.9 nm. Scale bar: 4 µm.

E. Confocal maximum intensity projections and optical sections of secondary spermatocytes during second meiotic division expressing indicated proteins. Z-size: 125.9 nm. Scale bar: 4 µm.

F. Confocal micrographs of an isolated male-derived spermatocyte and budding spermatids showing subcellular distribution of both mitochondria and PEI-1_ΔBTB+BACK+IDR::mTagRFP-T. White arrow heads mark budding spermatids, red arrows and dashed circles indicate residual bodies. Hoechst 33342 was used to stain DNA. MitoTracker® Green FM was used to stain mitochondria. Scale bars: 4 µm.

G. Line profiles display fluorescence intensity for PEI-1_ΔBTB+BACK+IDR::mTagRFP-T (magenta) versus mitochondria (green) signals over indicated line (shown in F). Vertical lines and colored circles indicate fluorescence peaks. a.u. – arbitrary unit.

**Figure S5. Related to Figure 5.**

A-E. Amino acid composition profiles of the intrinsically disordered region of PEI-1 (A), PGL-1(B), PGL-3 (C), MEG-3(D) and MEG-4(E). Bars representing serine and glutamine are highlighted in green, glycine in blue. The profiles were generated using Composition profiler (Vacic et al., 2007). Sequences were analyzed against the SwissProt database. Statistical significance was tested using the two-sample t-test (***: $p \leq 0.001$, **: $p \leq 0.01$, *: $p \leq 0.05$).

**Figure S6. Related to Figure 6 and Discussion.**

A. Confocal micrographs of an isolated male-derived spermatocyte and budding spermatids showing subcellular distribution of both mitochondria and full-length PEI-1::mTagRFP-T. White arrow heads mark budding spermatids, red arrows and dashed circles indicate residual bodies. Hoechst 33342 was used to stain DNA. MitoTracker® Green FM was used to stain mitochondria. Scale bars: 4 µm.

B. Confocal micrographs of isolated spermatids expressing GFP::3xFLAG::WAGO-3 and SPE-45::mCherry. Scale bar: 4 µm.

C. Schematic representing protein domains and predicted S-palmitoylation sites of PEI-1. S-palmitoylation sites were predicted using CSS-Palm (v4.0) and GPS-Lipid (v1.0) (Ren et al., 2008; Xie et al., 2016).

**Figure S7. Related to Discussion.**

A. Phylogenetic analysis showing PEI-1 conservation within the *Caenorhabditis* genus. The phylogenetic three was generated using EggNOG (v4.5.1) (Huerta-Cepas et al., 2016). PEI-1 was defined as query and compared to all eukaryote entries.

B. Protein length and domain composition of six human BTB domain-containing proteins that resemble PEI-1 protein composition.

**Supplemental movie legends**

**Movie S1. Related to Figure 5.**

Time sequence of an isolated male-derived spermatocyte expressing GFP::3xFLAG::WAGO-3 as shown in Figure 5D. Images are confocal maximum intensity projections. Scale bar: 4 µm.

**Movie S2. Related to Figure 6.**

Confocal Z-stack of an isolated male-derived spermatocyte expressing PEI-1::mTagRFP-T as shown in Figure 6B. Hoechst 33342 was used to stain DNA. MitoTracker® Green FM was used to stain mitochondria. Scale bar: 4 µm.

**Movie S3. Related to Figure 6.**

Confocal Z-stack of isolated budding spermatids (male-derived) expressing PEI-1::mTagRFP-T as shown in Figure 6B. Hoechst 33342 was used to stain DNA. MitoTracker® Green FM was used to stain mitochondria. Scale bar: 4 µm.

**Movie S4. Related to Figure 6.**

Time sequence of an isolated male-derived spermatocyte expressing GFP::3xFLAG::WAGO-3 in absence of SPE-10. Images are confocal maximum intensity projections. Scale bar: 4 µm.

### Figure S1

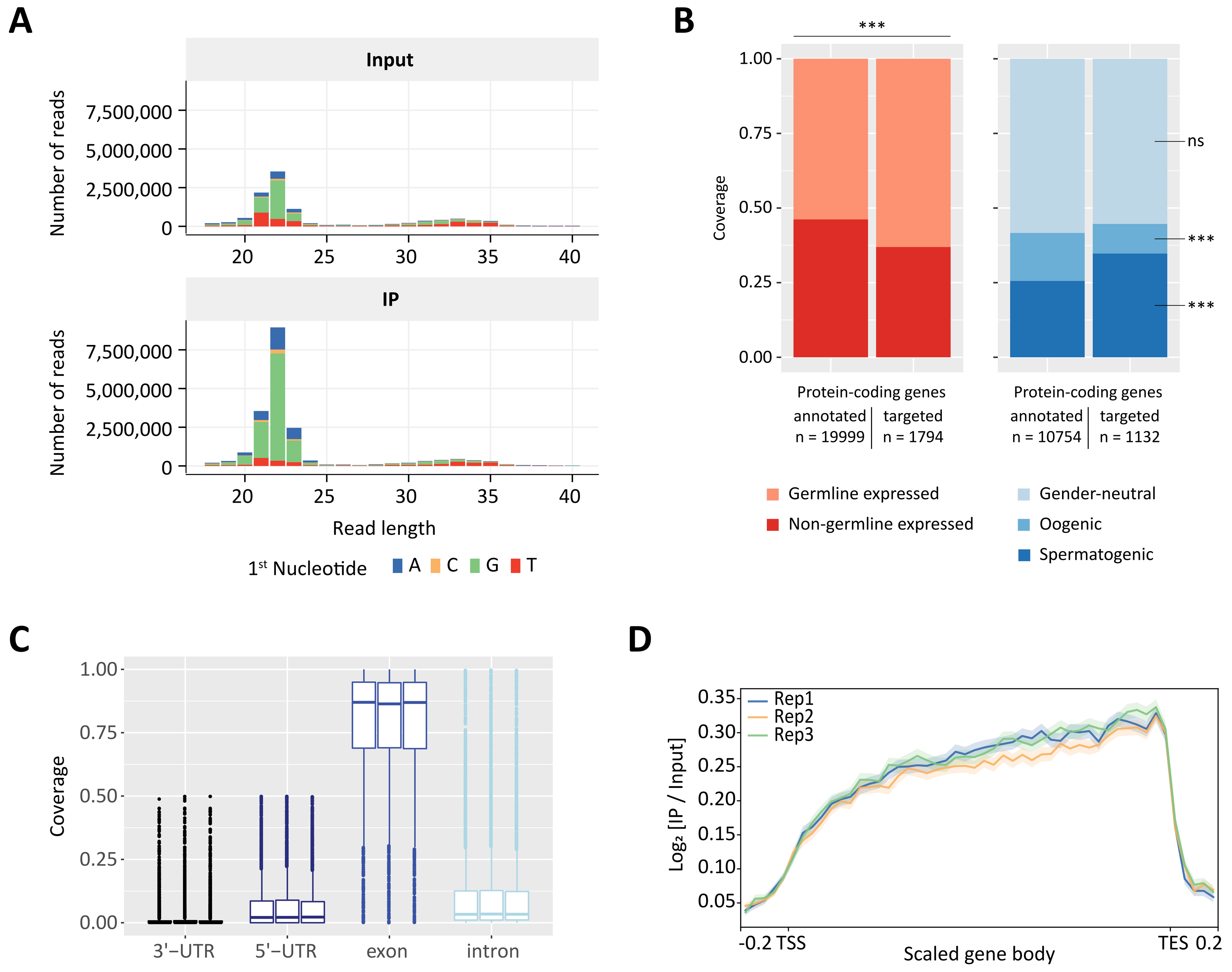
A
Input
IP
Number of reads
7,500,000
5,000,000
2,500,000
0
20
25
30
35
40
Read length
1st Nucleotide
A
C
G
T
B
***
1.00
0.75
0.50
0.25
0.00
Coverage
ns
Protein-coding genes
annotated
n = 19999
targeted
n = 1794
annotated
n = 10754
targeted
n = 1132
Germline expressed
Non-germline expressed
Gender-neutral
Oogenic
Spermatogenic
C
3'−UTR
5'−UTR
exon
intron
D
Rep1
Rep2
Rep3
Log2 [IP / Input]
0.35
0.30
0.25
0.20
0.15
0.10
0.05
-0.2 TSS
Scaled gene body
TES 0.2

### Figure S2

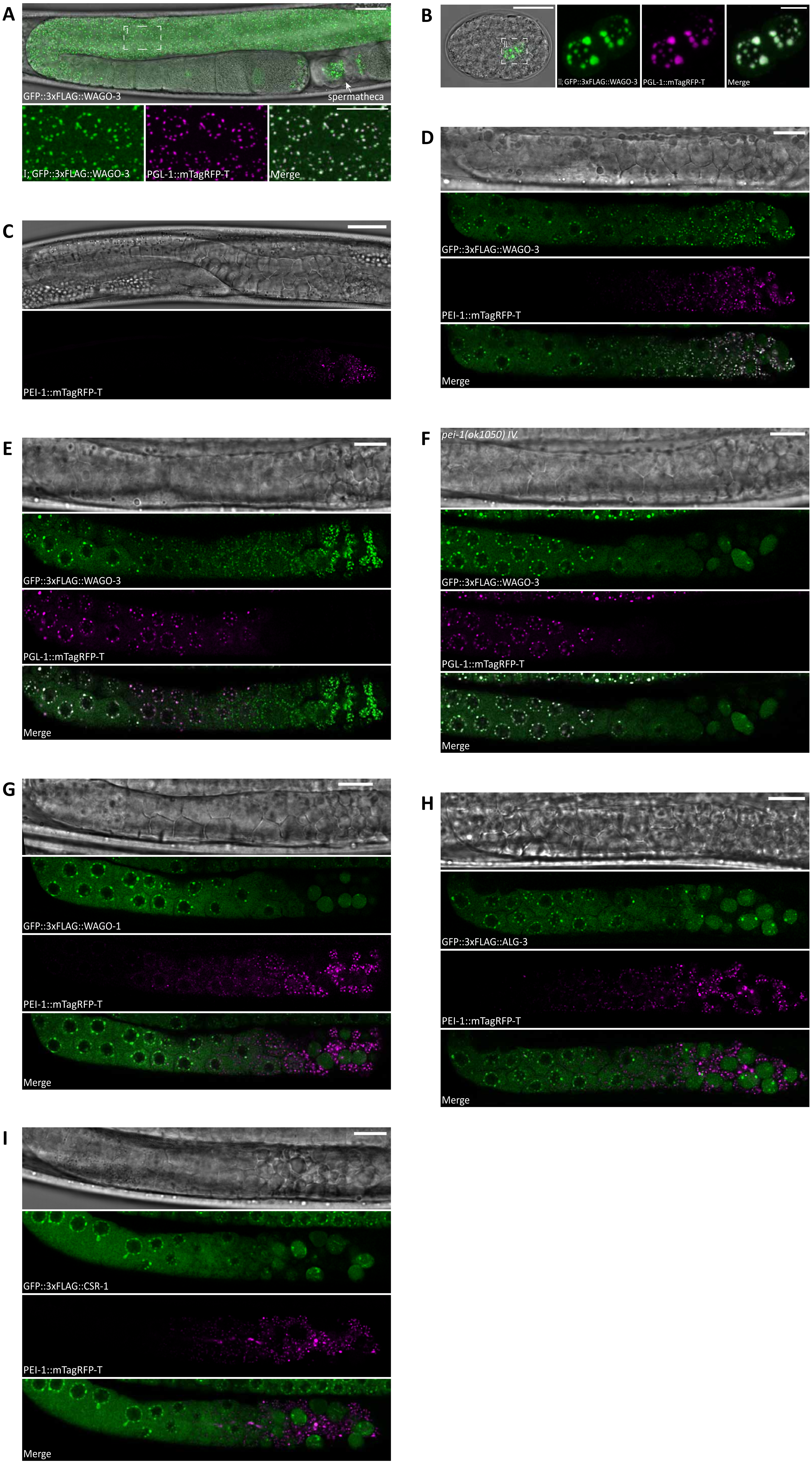
A
GFP::3xFLAG::WAGO-3
spermatheca
II; GFP::3xFLAG::WAGO-3
PGL-1::mTagRFP-T
Merge
B
II; GFP::3xFLAG::WAGO-3
PGL-1::mTagRFP-T
Merge
C
PEI-1::mTagRFP-T
D
GFP::3xFLAG::WAGO-3
PEI-1::mTagRFP-T
Merge
E
GFP::3xFLAG::WAGO-3
PGL-1::mTagRFP-T
Merge
F
pei-1(ok1050) IV.
GFP::3xFLAG::WAGO-3
PGL-1::mTagRFP-T
Merge
G
GFP::3xFLAG::WAGO-1
PEI-1::mTagRFP-T
Merge
H
GFP::3xFLAG::ALG-3
PEI-1::mTagRFP-T
Merge
I
GFP::3xFLAG::CSR-1
PEI-1::mTagRFP-T
Merge

### Figure S3

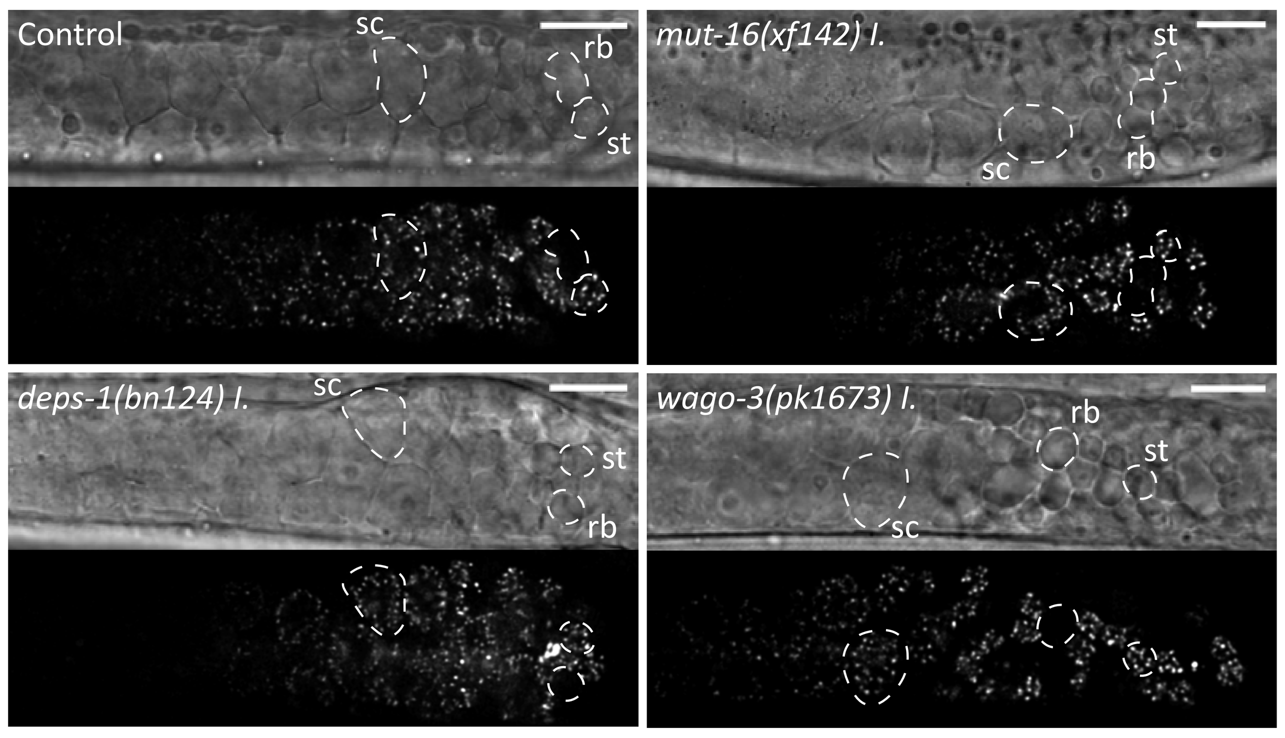

Control
sc
rb
st
*mut-16(xf142) I.*
st
sc
rb
*deps-1(bn124) I.*
sc
st
rb
*wago-3(pk1673) I.*
rb
st
sc

### Figure S4

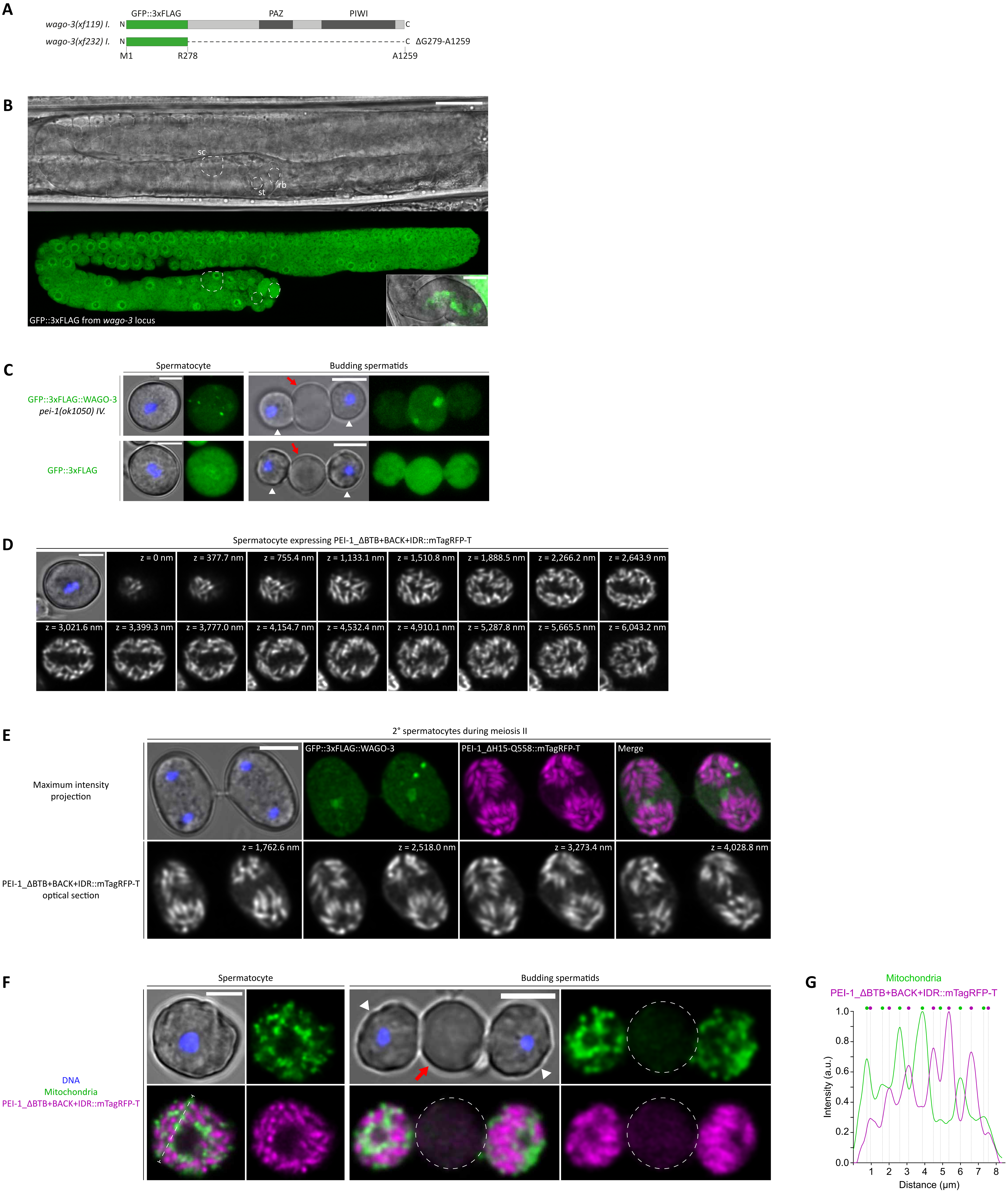
A
GFP::3xFLAG
PAZ
PIWI
wago-3(xf119) I.
wago-3(xf232) I.
N
C
ΔG279-A1259
M1
R278
A1259
B
sc
st
rb
GFP::3xFLAG from wago-3 locus
C
Spermatocyte
Budding spermatids
GFP::3xFLAG::WAGO-3
pei-1(ok1050) IV.
GFP::3xFLAG
D
Spermatocyte expressing PEI-1_ΔBTB+BACK+IDR::mTagRFP-T
z = 0 nm
z = 377.7 nm
z = 755.4 nm
z = 1,133.1 nm
z = 1,510.8 nm
z = 1,888.5 nm
z = 2,266.2 nm
z = 2,643.9 nm
z = 3,021.6 nm
z = 3,399.3 nm
z = 3,777.0 nm
z = 4,154.7 nm
z = 4,532.4 nm
z = 4,910.1 nm
z = 5,287.8 nm
z = 5,665.5 nm
z = 6,043.2 nm
E
2° spermatocytes during meiosis II
Maximum intensity projection
GFP::3xFLAG::WAGO-3
PEI-1_ΔH15-Q558::mTagRFP-T
Merge
PEI-1_ΔBTB+BACK+IDR::mTagRFP-T optical section
z = 1,762.6 nm
z = 2,518.0 nm
z = 3,273.4 nm
z = 4,028.8 nm
F
Spermatocyte
Budding spermatids
DNA
Mitochondria
PEI-1_ΔBTB+BACK+IDR::mTagRFP-T
G
Mitochondria
PEI-1_ΔBTB+BACK+IDR::mTagRFP-T
Intensity (a.u.)
Distance (μm)
0.0
0.2
0.4
0.6
0.8
1.0
1
2
3
4
5
6
7
8

### Figure S5

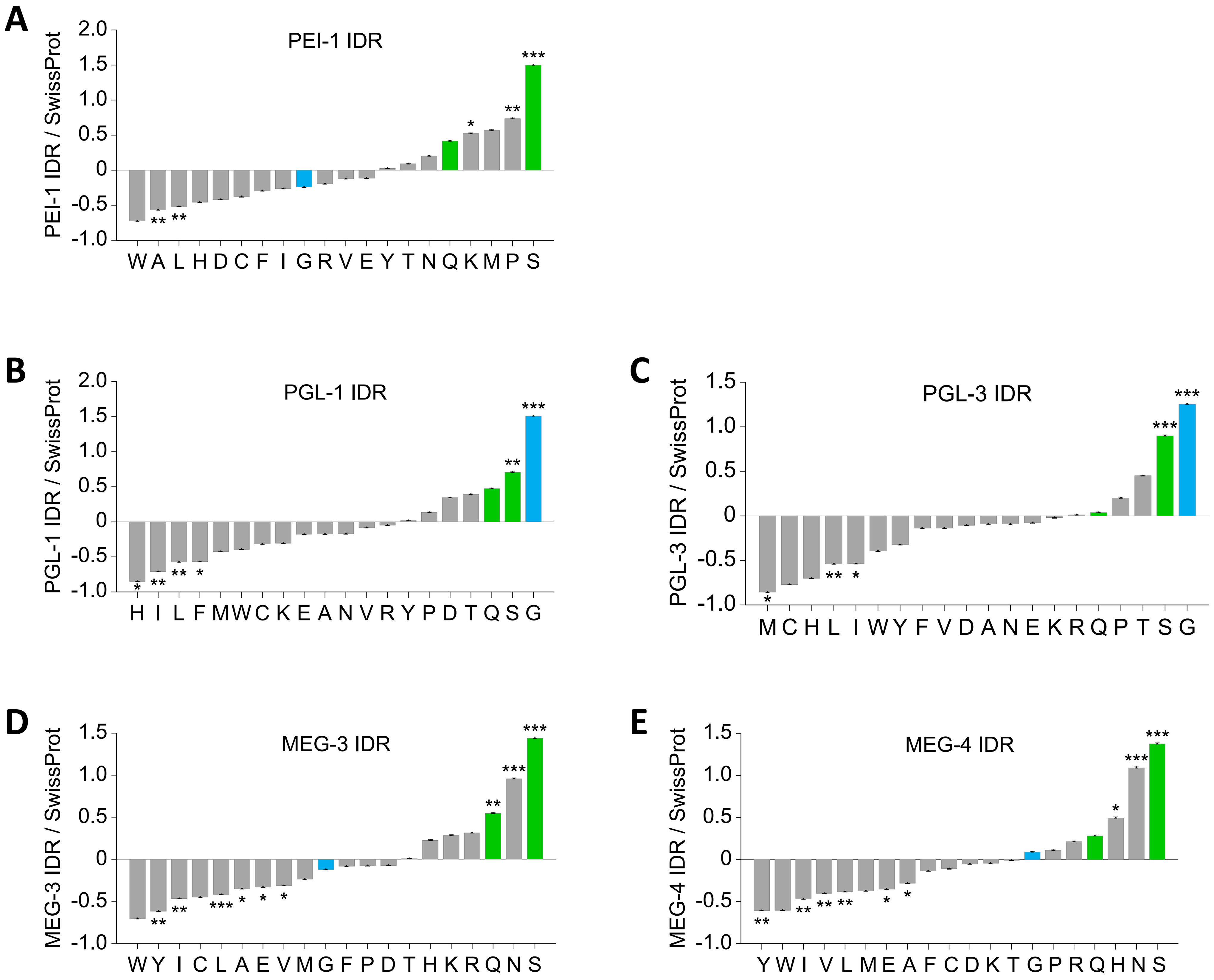

A
PEI-1 IDR
PEI-1 IDR / SwissProt
2.0
1.5
1.0
0.5
0
-0.5
-1.0
W A L H D C F I G R V E Y T N Q K M P S
*
**
**
*
**
***
B
PGL-1 IDR
PGL-1 IDR / SwissProt
2.0
1.5
1.0
0.5
0
-0.5
-1.0
H I L F M W C K E A N V R Y P D T Q S G
*
**
**
*
**
***
C
PGL-3 IDR
PGL-3 IDR / SwissProt
1.5
1.0
0.5
0
-0.5
-1.0
M C H L I W Y F V D A N E K R Q P T S G
*
**
*
***
***
D
MEG-3 IDR
MEG-3 IDR / SwissProt
1.5
1.0
0.5
0
-0.5
-1.0
W Y I C L A E V M G F P D T H K R Q N S
**
**
***
*
*
*
**
***
***
E
MEG-4 IDR
MEG-4 IDR / SwissProt
1.5
1.0
0.5
0
-0.5
-1.0
Y W I V L M E A F C D K T G P R Q H N S
**
**
**
**
*
*
*
***
***

### Figure S6

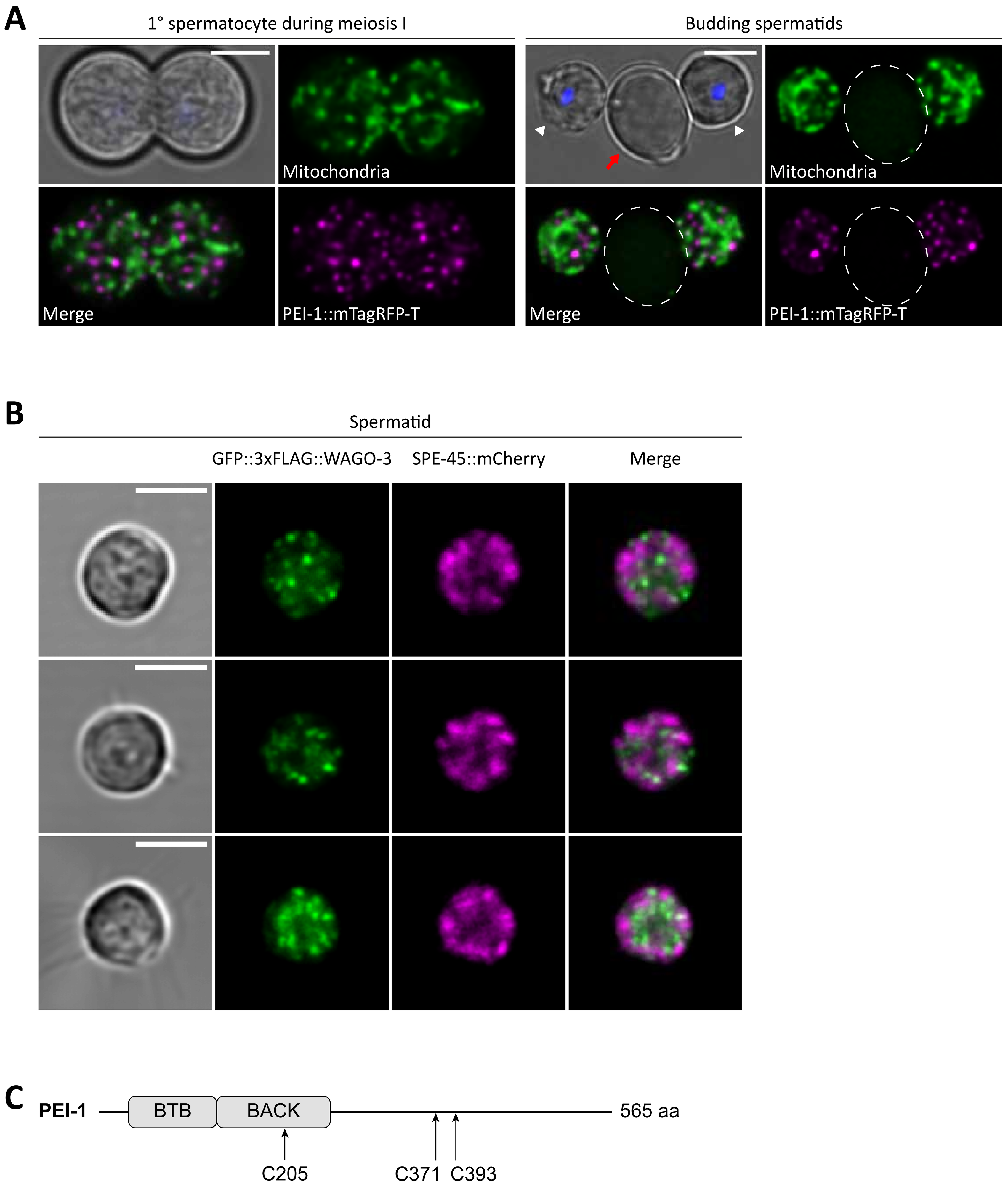
A
1° spermatocyte during meiosis I
Mitochondria
Merge
PEI-1::mTagRFP-T
Budding spermatids
Mitochondria
Merge
PEI-1::mTagRFP-T
B
Spermatid
GFP::3xFLAG::WAGO-3
SPE-45::mCherry
Merge
C
PEI-1
BTB
BACK
565 aa
C205
C371
C393

### Figure S7

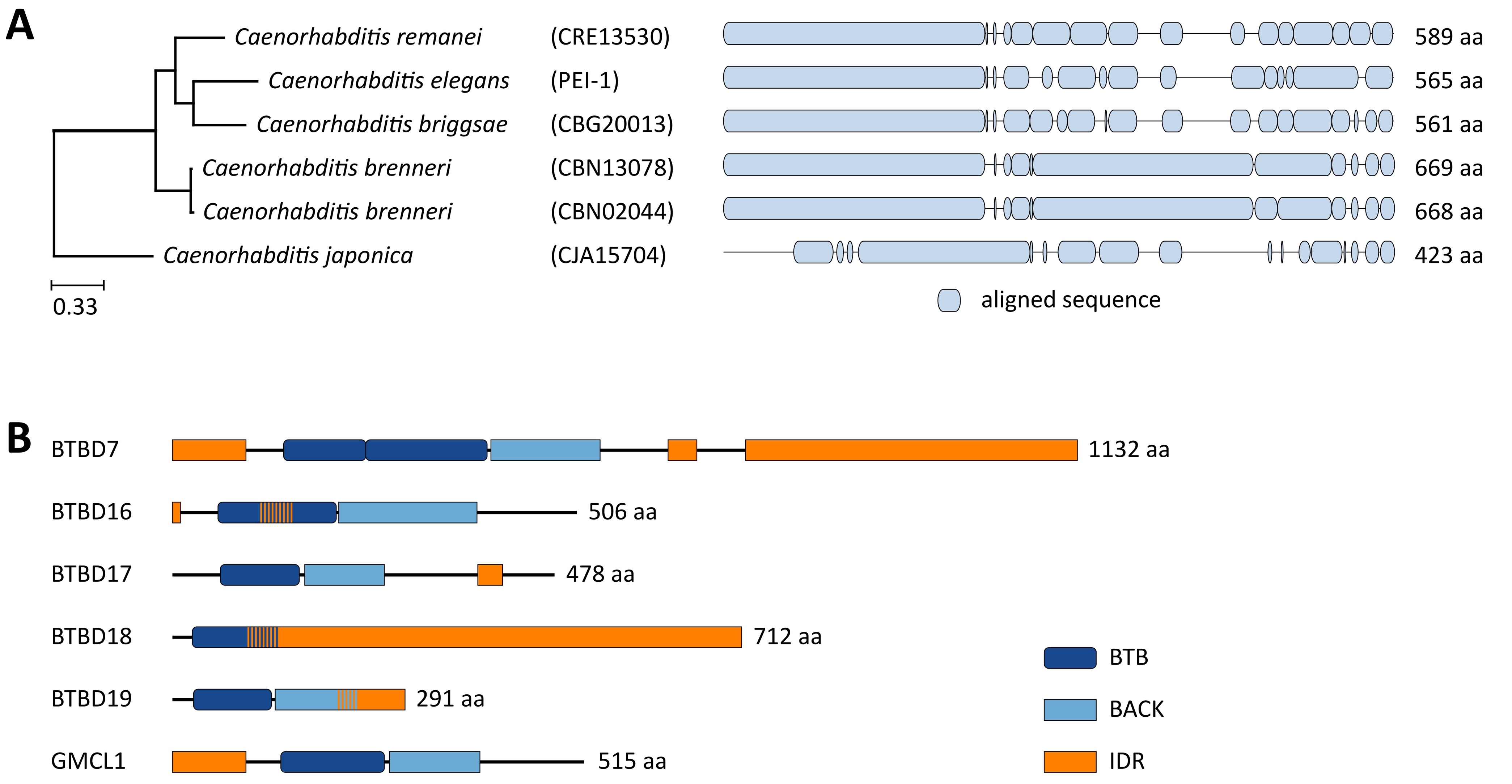
A
Caenorhabditis remanei
(CRE13530)
589 aa
Caenorhabditis elegans
(PEI-1)
565 aa
Caenorhabditis briggsae
(CBG20013)
561 aa
Caenorhabditis brenneri
(CBN13078)
669 aa
Caenorhabditis brenneri
(CBN02044)
668 aa
Caenorhabditis japonica
(CJA15704)
423 aa
0.33
aligned sequence
B
BTBD7
1132 aa
BTBD16
506 aa
BTBD17
478 aa
BTBD18
712 aa
BTBD19
291 aa
GMCL1
515 aa
BTB
BACK
IDR
